# Enzymatic Glycosylation of Anthranilates for Enhanced Functionality

**DOI:** 10.1101/2025.05.29.656899

**Authors:** Hani Gharabli, Allison M. Kohler, Carlotta Chiesa, Shreyas Wagle, Felipe Mejia-Otalvaro, Jean Victor Orth, Gonzalo Nahuel Bidart, Ann Dorrit Enevoldsen, Jochen Förster, Scott J. Werner, Ditte Hededam Welner

## Abstract

Anthranilate (ANT) is a precursor for the synthesis of valuable compounds, including its alkyl esters (AEANTs), such as methyl anthranilate (MANT). These derivatives are industrial petrochemical products used as flavouring agents and bird repellents. Due to the mandatory green transition, their biological industrial production must be considered. However, their antimicrobial activity and physicochemical properties inhibit efficient microbial production and challenge their practical use. To overcome this, we explored enzymatic glycosylation using UDP-dependent glycosyltransferases (UGTs). Screening identified three UGTs with activity on a selected AEANT panel, with UGT72B68 from *Solanum lycopersicum* showing the highest efficiency (840 s^−1^ M^−1^) for MANT. Rational engineering produced a mutant (F145M) with improved activity for bulkier AEANTs.

We scaled up enzymatic synthesis, producing 9.3 g of MANT-*N*-glucose (>99% purity, 74% yield). With this in hand, we observed that MANT-*N*-glucose has a significantly lower impact on the growth of *E. coli* and *P. putida*, supporting microbial production. Furthermore, we found that MANT-*N*-glucose completely inhibited sunflower seed consumption, compared to a 70% reduction observed in a previous study using MANT, when tested on captured red-winged blackbirds. Finally, a preliminary life-cycle assessment demonstrated that microbially produced MANT-*N*-glucose is a viable alternative to chemically synthesised MANT as a bird repellent.

## Introduction

Anthranilate (ANT) is a naturally occurring compound that is a key building block in the biosynthesis of tryptophan and various alkaloids.^**[1,2]**^ In addition to its biological significance, ANT plays an essential role in the chemical industry as a precursor to several commercial compounds, including alkyl esters of ANT (AEANTs), which are versatile in their applications.^**[3,4]**^ A notable example is methyl anthranilate (MANT), a naturally occurring compound responsible for Concord grapes’ distinctive scent and flavour.^**[5]**^ Hence, MANT is a flavour and fragrance compound used in the food and cosmetics sectors.^**[6]**^ Additionally, MANT is used as a bird and insect repellent,^**[7,8]**^ and as an antimicrobial agent against fungi and bacteria.^**[9,10]**^

There is a global market for ANT and its derivatives, which demands environmentally mild production methods to enable a sustainable chemical industry.^**[11]**^ Currently, the production of these compounds relies on petrochemicals, acid catalysts, and extreme reaction conditions.^**[4,12,13]**^ Alternative production methods for MANT, including the *N*-demethylation of methyl *N*-methyl anthranilate^**[14–16]**^ and the esterification of anthranilate^**[17,18]**^ through enzymatic or whole-cell biotransformation, have been investigated but found to be economically unfeasible.^**[14]**^ Consequently, researchers have developed ANT-and MANT-producing microbial cell factories.^**[11,19,20]**^ However, the achievable titers are limited by the host toxicity of these compounds.^**[3,20]**^ In addition, MANT is volatile and poorly soluble in water, which, along with its phytotoxic activity,^**[21,22]**^ challenges the formulation and utility of MANT for agricultural applications.^**[23]**^ Other AEANTs, such as ethyl and butyl anthranilate (EANT and BANT, respectively), display similar activities as MANT but exhibit even lower water solubility.^**[24]**^

In nature, many of the adverse properties presented by AEANTs are alleviated through a glycosylation step.^**[25,26]**^ Glycosylation often has a significant effect on the physicochemical properties of the compound, including solubility and volatility,^**[27]**^ and can be used as a route for lower toxicity,^**[28,29]**^ increased stability,^**[30]**^ and an altered flavour profile.^**[31]**^ In nature, glycosylation reactions are mainly catalysed by glycosyltransferases (GTs). Among these, the family 1 GTs are promising substitutes for conventional chemical glycosylation to produce valuable natural glycosides or ‘new-to-nature’ glycosides.^**[32,33]**^ This is owed to their acceptor substrate promiscuity, efficient one-step regio-and stereo-selective glycosylation reaction, and mild reaction conditions.^**[34]**^ They utilise a nucleotide-activated sugar donor to transfer a sugar moiety to an acceptor substrate, most often uridine diphosphate (UDP) glucose,^**[35]**^ and are commonly denoted as UDP-dependent glycosyltransferases (UGTs). While no UGT has been identified with activity on the naturally occurring AEANTs, UGTs with activity towards ANT have been discovered.^**[36–38]**^ In addition, UGTs responsible for glycosylating methyl salicylate, a close homologue of MANT harbouring a phenol in the corresponding position of the aniline, have also been identified,^**[39,40]**^ making it probable that UGTs with AEANT activity exist in nature along with the resulting *N*-glycosides.

Given that glycosylation has the potential to mitigate many of the adverse properties of AEANTs, our objective was to discover and characterise UGTs capable of glycosylating a panel of safe-to-consume AEANTs. This included MANT, EANT, BANT, isobutyl anthranilate (IBANT), and cyclohexyl anthranilate (CANT). By screening an in-house UGT library, we identified three UGTs—UGT71C1, UGT72B19, and UGT72B68—that could efficiently glycosylate multiple AEANTs. Among the three identified UGTs, UGT72B68 exhibited the highest catalytic efficiency with MANT, making it the primary focus of this study. This included the rational engineering of its active site to broaden its AEANT acceptor specificity along with its utilisation in the biocatalytic production and isolation of MANT-*N*-glucose on a multigram scale. The bird-repellent and antimicrobial activities of the isolated glucoside were subsequently tested. This provided data for a preliminary life cycle assessment (LCA), which elucidated the environmental consequences of substituting petrochemical MANT with MANT-*N*-glucose. The findings reported in this study highlight the potential of using biocatalytic glycosylation as a method for producing novel bioactive compounds with enhanced physicochemical properties and potential scalability through microbial production.

## Results and Discussion

### Discovery and Characterisation of Three AEANT-Glycosylating UGTs

An in-house library of plant UGTs[41] was employed to discover UGTs capable of glycosylating the selected panel of AEANTs (Figure 1a). Of the 19 UGTs tested with MANT, 9 showed more than 50 % conversion, while 8 of the 28 UGTs assayed with EANT exhibited more than 50 % conversion (Figure 1b, Supplementary Table 6). These UGTs were then tested with BANT, leading to the selection of three UGTs that demonstrated the highest overall activity across all three substrates. These were UGT71C1 from *Arabidopsis thaliana* (thale cress) (UniProt accession number O82381), UGT72B19 from *Fagopyrum esculentum* (buckwheat) (UniProt accession number A0A0A1H7P3), and UGT72B68 from *Solanum lycopersicum* (tomato) (UniProt accession number D7S016). These UGTs were subsequently assayed with IBANT and CANT. Here, UGT72B68 showed activity with IBANT but not with CANT. In contrast, UGT72B19 and UGT71C1 demonstrated activity with both acceptors, with UGT71C1 exhibiting the highest overall activity (Figure 1b, Supplementary Table 6). Representative chromatograms for each AEANT and their corresponding glucoside are shown in Figure 1c.

**Figure 1.**
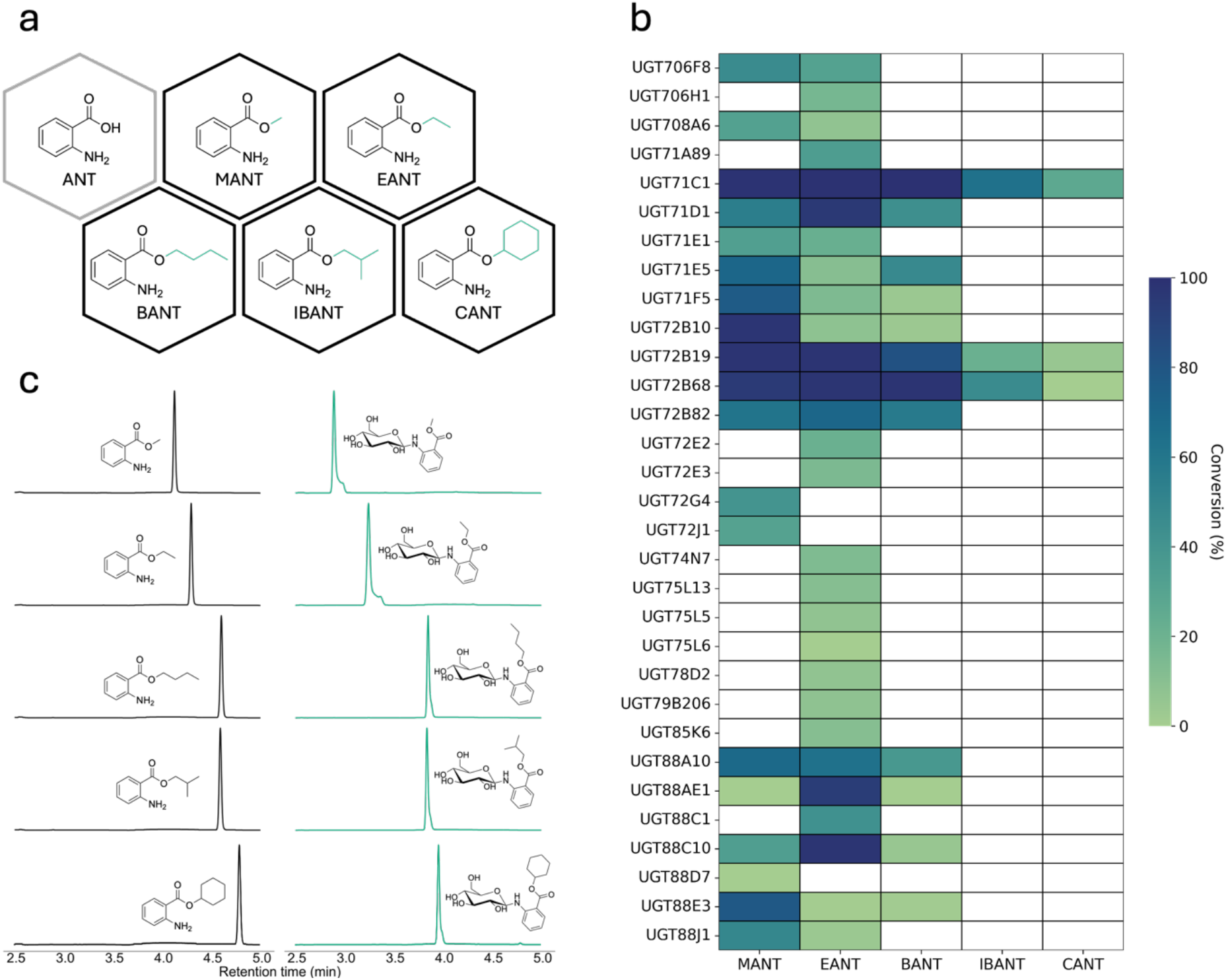
(a) The chemical structure of ANT and the panel of AEANTs tested in this study. (b) A heatmap displaying the conversion percentage achieved after an overnight reaction between UGT and the AEANT panel. A white panel indicates an enzyme-substrate pair that was not tested. (c) Chromatograms of the AEANTs (in black) and their corresponding glycoside (in teal) produced using WT or the F145M variant of UGT72B68.

UGT71C1 shares 33.8 % and 31.3 % sequence identity with UGT72B19 and UGT72B68, respectively, and UGT72B19 and UGT72B68 share 61.8 % sequence identity. UGT71C1 has previously been reported to glycosylate flavonols, such as kaempferol and quercetin.^[42,43]^ UGT72B19 and UGT72B68 have been reported to glycosylate 3,4-dichloroaniline.^[44]^ Interestingly, UGT72B68 has also been reported to accept methyl salicylate, which is structurally similar to MANT, as a substrate.^[40]^ However, MANT has not yet been identified in tomato, the plant from which UGT72B68 originates.

### Biochemical Characterisation of UGT71C1, UGT72B19, and UGT72B68

UGT71C1, UGT72B19, and UGT72B68 were biochemically characterised using MANT as the acceptor substrate and UDP-glucose as the donor substrate. First, the pH and temperature optima were determined (Supplementary Figures 2 and 3). UGT71C1 and UGT72B19 displayed optimal activity in phosphate buffer, pH 8.0 (Supplementary Figure 2). UGT72B68 exhibited similar activity in phosphate buffer (pH 8.0) and glycine buffer (pH 9.0). To streamline subsequent experiments, phosphate buffer (pH 8.0) was chosen. The temperature optima were found to be 30 °C for UGT71C1, 39 °C for UGT72B19, and 35 °C for UGT72B68 (Supplementary Figure 3). These reaction conditions were used to establish the kinetic properties of the UGTs (Table 1, Supplementary Figure 4). All UGTs exhibited Michaelis-Menten-like kinetics, with UGT72B68 having the highest catalytic efficiency with MANT. Based on this, UGT72B68 was selected for further investigation, beginning with establishing its kinetic properties using EANT, BANT, and IBANT as acceptor substrates (Table 1 and Supplementary Figure 5). UGT72B68 displayed the highest catalytic efficiency with BANT as the acceptor substrate and the lowest with MANT. Meanwhile, it displayed the highest maximum velocity (V_max_) with EANT, which, combined with the higher Michaelis constant (K_m_), could be related to the equilibrium between substrate binding and product release.^[45]^ This is further discussed in the following section.

**Table 1.** Kinetic properties of UGTs with MANT, EANT, BANT, and IBANT

| UGT isoform | Acceptor | $K_m$ ($\mu\text{M}$) | $V_{\max}$ ($\mu\text{M min}^{-1}$) | $k_{\text{cat}}$ ($\text{s}^{-1}$) | $k_{\text{cat}}/K_m$ ($\text{s}^{-1} \text{M}^{-1}$) |
| --- | --- | --- | --- | --- | --- |
| UGT71C1 | MANT | $164.5 \pm 22.0$ | $2.4 \pm 0.7$ | $0.079 \pm 0.022$ | 473 |
| UGT72B19 | MANT | $212.2 \pm 19.1$ | $2.8 \pm 0.9$ | $0.042 \pm 0.013$ | 203 |
| UGT72B68 | MANT | $347.1 \pm 45.7$ | $12.1 \pm 1.2$ | $0.287 \pm 0.027$ | 840 |
| UGT72B68 | EANT | $889.2 \pm 43.1$ | $38.8 \pm 3.7$ | $2.157 \pm 0.205$ | 2423 |
| UGT72B68 | BANT | $226.8 \pm 17.7$ | $15.6 \pm 0.8$ | $1.038 \pm 0.055$ | 4602 |
| UGT72B68 | IBANT | $259.9 \pm 47.6$ | $4.0 \pm 0.1$ | $0.267 \pm 0.008$ | 1048 |

### Identification of a UGT72B68 Variant with Broadened Specificity and Activity

Among the three identified AEANT-glycosylating UGTs, UGT71C1 exhibited the broadest selectivity, showing activity with CANT and slightly higher activity with IBANT than UGT72B68 and UGT72B19 (Figure 1b, Supplementary Table 6). To investigate this observation, we set out to engineer UGT72B68 to enable the glycosylation of CANT. Here, we hypothesised that this could be engineered by altering its active site residue composition. First, the approximate AEANT binding site of UGT72B68 was identified by docking MANT, EANT, BANT, IBANT, and CANT into UGT72B68, in which a common orientation was observed (Supplementary Figure 6). This could indicate that this orientation is the preferred binding mode of the AEANTs in UGT72B68. Using the UGT72B68/MANT complex structure, we identified the residues within 5 Å of MANT and compared them to the corresponding residues in UGT71C1 (Supplementary Figure 7).

A different distribution of aromatic residues was observed between the active sites of the two UGTs (Figure 2a, b), which was hypothesised to play a role in their respective specificities. To test this hypothesis, all positions in UGT72B68 occupied by an aromatic residue in either of the two enzymes were mutated to the corresponding residue in UGT71C1, generating five UGT72B68 single mutant variants (Figure 2c). In addition, variant I80A was tested due to the residue’s position relative to the alkyl-ester of the AEANTs (Figure 2a).

**Figure 2.**
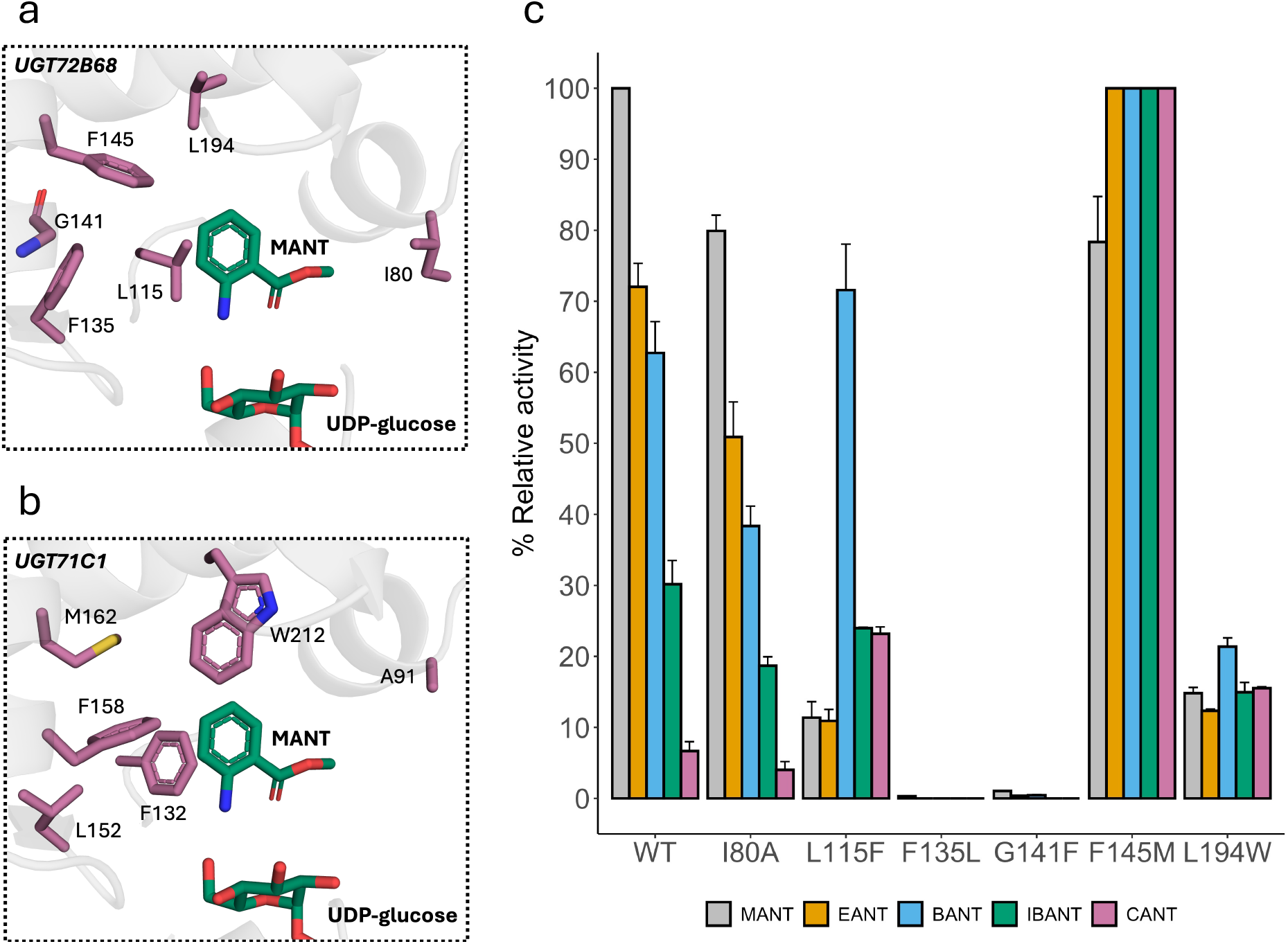
(a, b) Residues targeted for mutagenesis in UGT72B68 (a) and the corresponding residues in UGT71C1 (b) shown in stick representations along with MANT and UDP-glucose displayed in teal. (c) Relative activity of the UGT72B68 variants. Activity values were normalised to the variant with the highest activity for each substrate.

The activities of the resulting six UGT72B68 variants were tested against the AEANT panel (Figure 2c). We found that the I80A mutation has no impact on the AEANT selectivity of UGT72B68, while mutations F135L and G141F abolished activity. The loss of activity in F135L may result from the loss of an important pi-pi interaction, while other aromatic residues in UGT71C1 could fulfil this role, such as F158. In contrast, the introduction of a bulky aromatic residue in position 141 may sterically hinder substrate binding in UGT72B68, with two aromatic residues already occupying this space.

Interestingly, the L115F variant exhibited decreased activity with MANT and EANT, comparable activity for IBANT, but increased activity for BANT and CANT compared to the WT. While the underlying mechanism is unclear, the phenylalanine could form a T-shaped pi-pi interaction if oriented as observed in UGT71C1. This could stabilise substrate binding but may also stabilise the product, inhibiting its release. This could explain the lower K_m_ and k_cat_ observed in UGT71C1 compared to UGT72B68 with MANT. However, it is still unclear why this effect is only observed with three of the AEANTs.

Remarkably, variant F145M displayed significantly enhanced activity with all bulky AEANTs while still retaining 80 % WT activity with MANT. This increased activity could be caused by the smaller size and higher flexibility of methionine compared to phenylalanine, which could increase the flexibility and size of the binding cavity, facilitating the reactive binding of the bulkier AEANTs. The importance of this residue in the substrate specificity of UGTs has also been reported in a previous study.^**[46]**^ Here, the authors reported that substituting a glutamic acid with a range of residues, particularly with isoleucine or methionine, enabled the bis-*C*-glycosylation of 2-Phenyl-2′,4′,6′-trihydroxyacetophenone. However, this effect was also achieved via the introduction of phenylalanine, seemingly not being related to the effects observed in this study.

The recent emergence of accurate modelling of enzyme-substrate complexes offers an alternative to conventional molecular docking approaches.^**[47]**^ Taking advantage of readily available tools, the structural complexes of UGT72B68, the AEANTs, and UDP-glucose were prepared using Chai-1^**[48]**^ to validate our findings. Here, UDP-glucose was bound as observed in previously elucidated crystal structures across all predictions (Supplementary Figure 8),^**[49,50]**^ and a MANT binding pose similar to that predicted by molecular docking was found. However, the remaining AEANTs (EANT, BANT, IBANT, and CANT) were observed in similar binding coordinates as observed in the molecular docking poses but flipped 180 degrees along the vertical axis (Supplementary Figure 8). This results in the alkyl-ester being in proximity to F145 instead of I80, offering a rationalisation of the observed effects of F145 on AEANT specificity (Figure 2c).

### Multigram-scale production and purification of MANT-*N*-glucose

Having identified an efficient enzymatic MANT-*N*-glucose catalyst, and since we are interested in enhancing global sustainability, in which agricultural technology is a major player, we wanted to test the bird deterrence properties of MANT-*N*-glucose. To facilitate this, we needed the pure compound on a multigram scale. Therefore, we scaled up the biocatalytic conversion of MANT to MANT-*N*-glucose and developed a purification method. Sucrose synthase, particularly from *Glycine max* (*Gm*SuSy), can be used in combination with a UGT to regenerate UDP-glucose.^[51,52]^ However, in accordance with the literature, we found that *Gm*SuSy has low chemotolerance (Figure 3a) while also needing a high sucrose load, making it unsuitable for our multi-gram scale reaction.^[53,54]^ The chemotolerance of UGT72B68 towards MANT was also tested since we have previously observed the inactivation of UGTs at substrate loads in the millimolar range.^[55,56]^ UGT72B68 displayed remarkable tolerance to MANT concentrations up to 20 mM, which also required tolerance of up to 20% (v/v) DMSO, the solvent for the MANT stock solution (Figure 3b, c).

**Figure 3.**
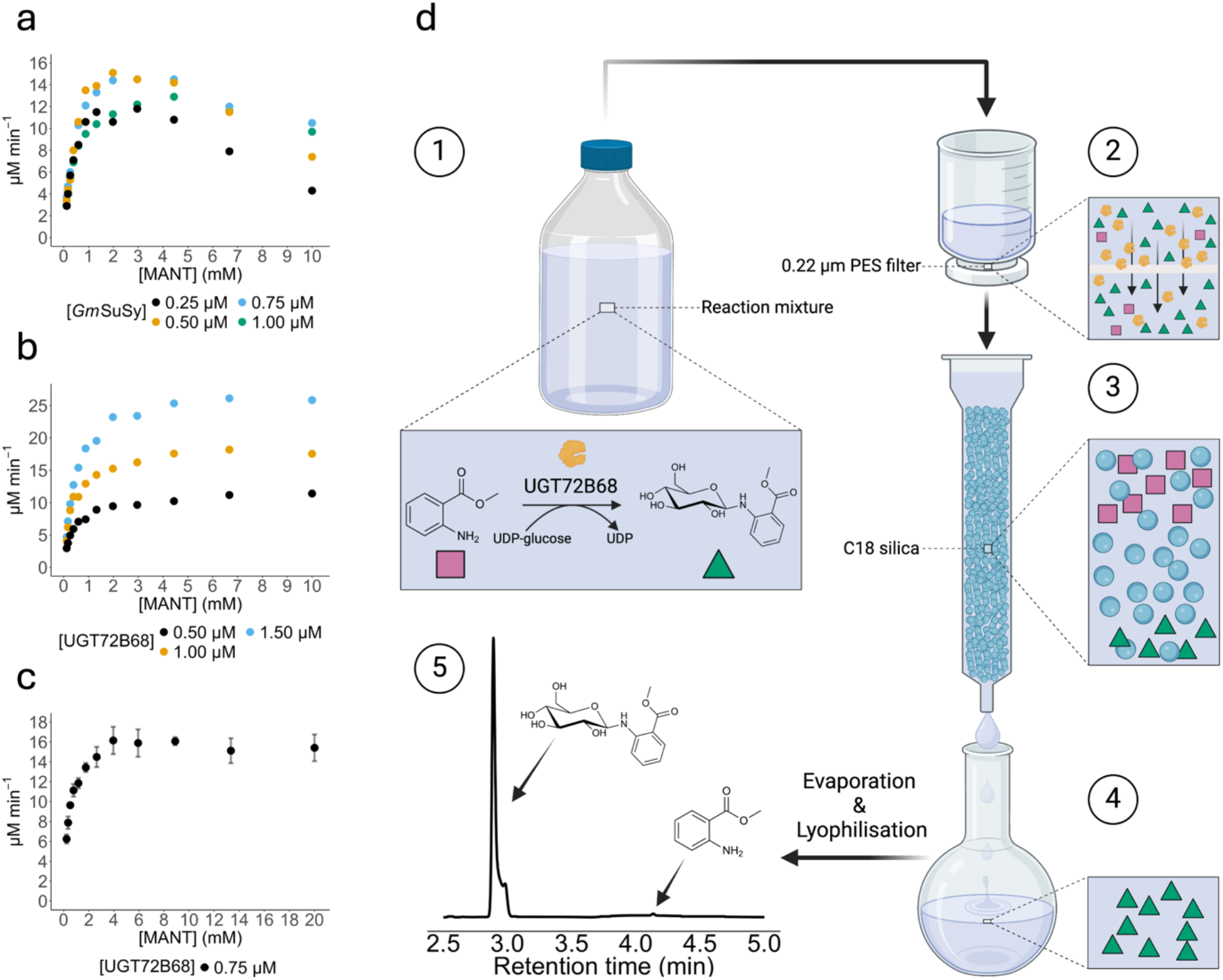
(a) The chemotolerance of GmSuSy towards MANT. (b) The chemotolerance of UGT72B68 towards MANT concentrations up to 10 mM. (c) The chemotolerance of 0.75 µM UGT72B68 against MANT concentrations up to 20 mM (d) An overview of the process for the gram-scale production of MANT-N-glucose. (d1) The reaction was performed in a flask containing 20 mM MANT, 30 mM UDP-glucose, and 221 mg UGT72B68, which was left to react for 16 hours. (d2) The reaction mixture was subsequently filtered through a 0.22 µm PES sterile filter to remove the denatured enzymes and any formed solids from the mixture. (d3) The product was then separated from the other constituents via flash chromatography, employing a column containing C18 silica particles, including the remaining enzymes in the mixture. (d4) The fractions containing the product were collected, pooled and then evaporated in vacuo and subsequently lyophilized to remove residual moisture. (d5) The final product was achieved at > 99 % purity, as determined by HPLC.

UGT72B68 was purified from the lysate of 22 L *E. coli* culture. This yielded 221 mg purified UGT72B68, which we utilised in a 2 L reaction consisting of 20 mM MANT and 30 mM UDP-glucose (Figure 3d). After overnight incubation, 90 % of MANT was converted to MANT-*N*-glucose (Supplementary Figure 9). The separation of the product from the remaining constituents of the reaction mixture was achieved via flash chromatography. Subsequent evaporation *in vacuo* and lyophilisation yielded 9.3 g of an off-white powder at > 99 % purity (HPLC)

(Figure 3d). 1D and 2D NMR confirmed the powder to be MANT-*N*-glucose (Supplementary Figures 10-12). The resulting product yield was 74.4 % from the reaction, with a recovery rate from the purification process of 82.3 %. The biocatalytic conversion of MANT to MANT-*N*-glucose can reach 100 % conversion, as observed in the initial screening (Figure 1b, Supplementary Table 6). However, this was not achieved, which could be caused by the hydrolysis of UDP-glucose, the denaturation of UGT72B68 over time, or insufficient reaction time. The reaction could potentially be improved by spiking with fresh enzyme, acceptor substrate, and UDP-glucose to achieve a higher conversion yield and increase the final concentration of the product in the reaction mixture.

### Antimicrobial Effects of MANT-*N*-Glucose

As mentioned, previously reported cell factories for MANT developed to replace petroleum-based synthesis are limited by MANT’s toxicity towards the microbial host.^[20]^ To investigate if this effect is alleviated by glycosylation, the growth of two common cell factory strains, *E. coli* and *P. putida*, was monitored upon exposure to a range of MANT and MANT-*N*-glucose concentrations up to 7.5 mM (Figure 4a-d). In accordance with a previous study,^[20]^ both organisms exhibited a dose-dependent growth inhibition by MANT, with *P. putida* seemingly being more sensitive than *E. coli* (Figure 4a, b). However, no response to MANT-*N*-glucose was found in either of the organisms when subjected to the same conditions as with MANT (Figure 4c, d). We then probed the two strains with production-relevant MANT-*N*-glucose concentrations (up to 50 mM), which revealed a dose-dependent inhibition of *E. coli*. However, no change in the growth of *P. putida* was observed. This indicates that glycosylation could serve as a strategy to enhance cell factory titres of MANT and other AEANTs, with *P. putida* being a good candidate for a host strain.

**Figure 4.**
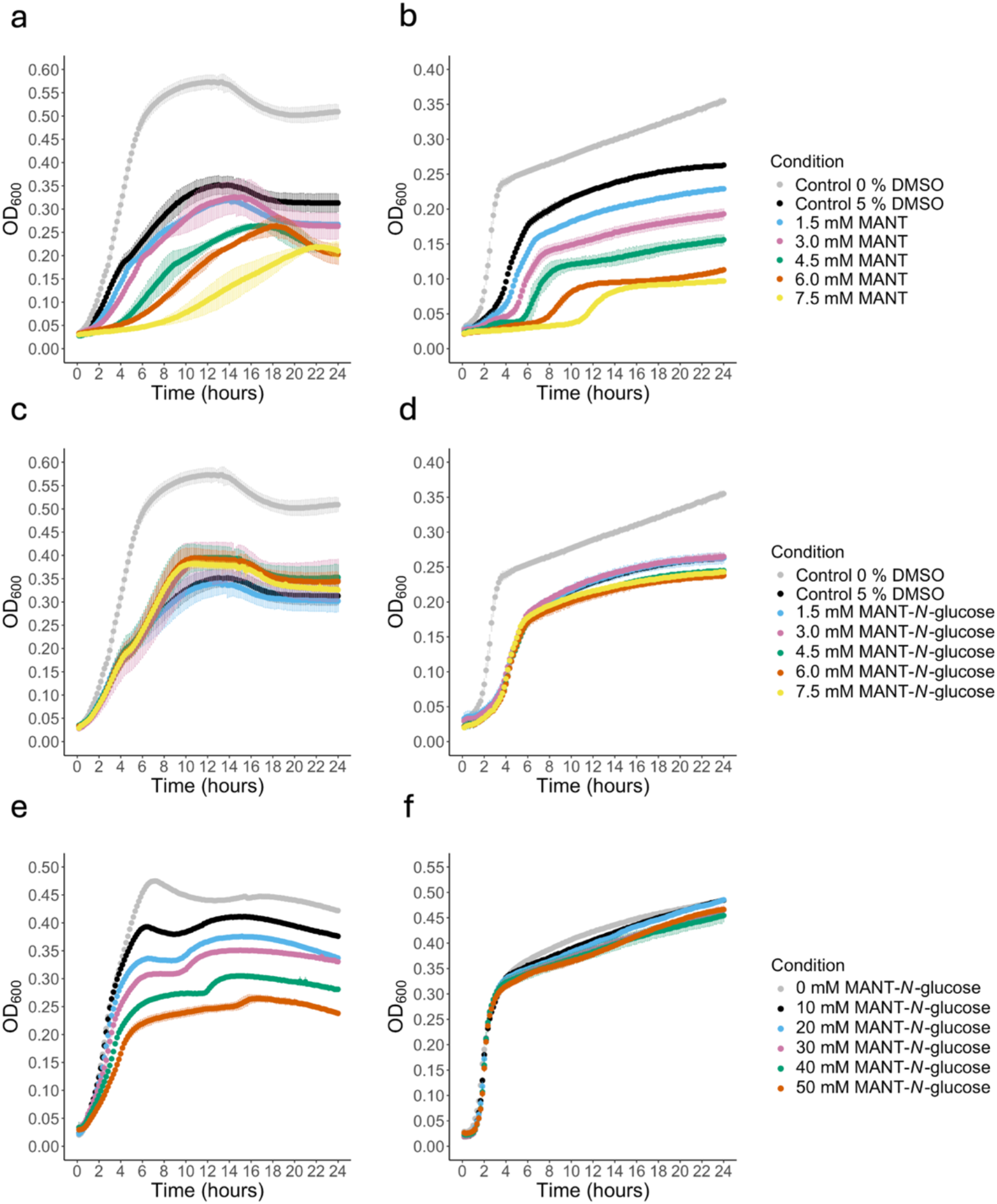
The effect on growth of MANT on E. coli (a) and P. putida (b) and MANT-N-glucose on E. coli (c) and P. putida (d). A control of 5% DMSO (the maximum, added in the 7.5 mM samples) was included. (e) and (f) display the growth of E. coli and P. putida at higher MANT-*N*-glucose concentrations. No DMSO is needed with MANT-*N*-glucose and was, therefore, omitted in these two experiments (e, f).

### Chemical Degradation of MANT-*N*-Glucose

After demonstrating that MANT-*N*-glucose exhibits limited antimicrobial activity (Figure 4), the biobased production of MANT-*N*-glucose remains a feasible strategy. However, the desired product may still be the corresponding aglycone, MANT. Therefore, we examined the possibilities of retrieving the aglycon from the corresponding *N*-glucoside via acid-and base-catalysed hydrolysis. We found that 100 mM HCl cleaves the glycosidic bond of 5 mM MANT-*N*-glucoside to completion within an hour (Supplementary Figure 13). Complete cleavage was also achieved with 10 mM HCl, but only within approximately 5 hours. These results correspond well with previous observations.^[57]^ When using a base, the *N*-glycosidic bond was stable. Instead, the ester was cleaved to reveal the free acid (Supplementary Figure 14). This was validated via LC-MS (Supplementary Figure 15). Interestingly, this glycosidic compound (ANT-*N*-glucose) can likely not be achieved via UGT-mediated glycosylation of ANT, since the enzyme would target the carboxylic acid with a low product yield (data not shown).

### Bird Repellence of MANT-*N*-Glucose

Anthranilates, including MANT, are widely implemented as benign bird control agents, acting by repelling the birds from an area without harming them.^[8]^ Since glycosylation changes the physicochemical properties of a compound, including its volatility and solubility, the bird deterrence of MANT-*N*-glucose could be drastically changed compared to MANT. With the pure MANT-*N*-glucose in hand, we tested its bird repellence in a one-day two-choice test with MANT-*N*-glucose-treated and untreated sunflower seeds on red-winged blackbirds in captivity (Supplementary Table 1). The MANT-*N*-glucose was dosed at 1.0 % (w/w). The blackbirds consumed significantly more untreated sunflower during the preference experiment (P = 0.0005). During the experiment, blackbirds consumed an average of 0 ± 0.7 g of treated sunflower and 6.9 ± 0.8 g of untreated sunflower, indicating the retained bird-repellent properties of the glycoside. Interestingly, in a previous study using the same experimental setup and treatment dose but with rice as the feed, the blackbirds consumed an average of 0.8 ± 0.2 g of MANT-treated rice compared to 2.6 ± 0.3 g of untreated rice (70% repellency).^[58]^ In contrast, we saw no consumption of glycoside-treated seeds (100% repellency), indicating an improved efficacy of the glycoside compared to the aglycon. However, further studies should be conducted to validate this, including a dose-dependency experiment to understand the optimal MANT-*N*-glucose concentration.

### The environmental impacts of substituting MANT with MANT-*N*-glucose as a bird repellent

Having shown that MANT-*N*-glucose is a potent bird repellent agent, we wondered if the potential environmental advantage of substituting MANT for MANT-*N*-glucose outweighs the footprint from the additional glycosylation reaction. This was assessed through a preliminary comparative LCA, with the production of 100 g of sunflower achenes as the functional unit. The environmental impacts were evaluated across three scenarios (Supplementary Figure 1). The scenarios included chemical synthesis of MANT (scenario 1), chemoenzymatic synthesis of MANT-*N*-glucose (scenario 2), and microbial synthesis of MANT-*N*-glucose (scenario 3). Despite its greater bird repellence, chemoenzymatic MANT-*N*-glucose showed higher impacts in all midpoint categories except for land use (LU) and water consumption (WC) when compared to scenario 1 (Supplementary Table 7). At the end-point level, chemoenzymatic MANT-*N*-glucose demonstrated a 10 % reduction in ecosystem quality impact but increases of 62 % and 73 % in human toxicity impact and resource scarcity impact, respectively, compared to scenario 1 (Figure 5a, b, and c).

**Figure 5.**
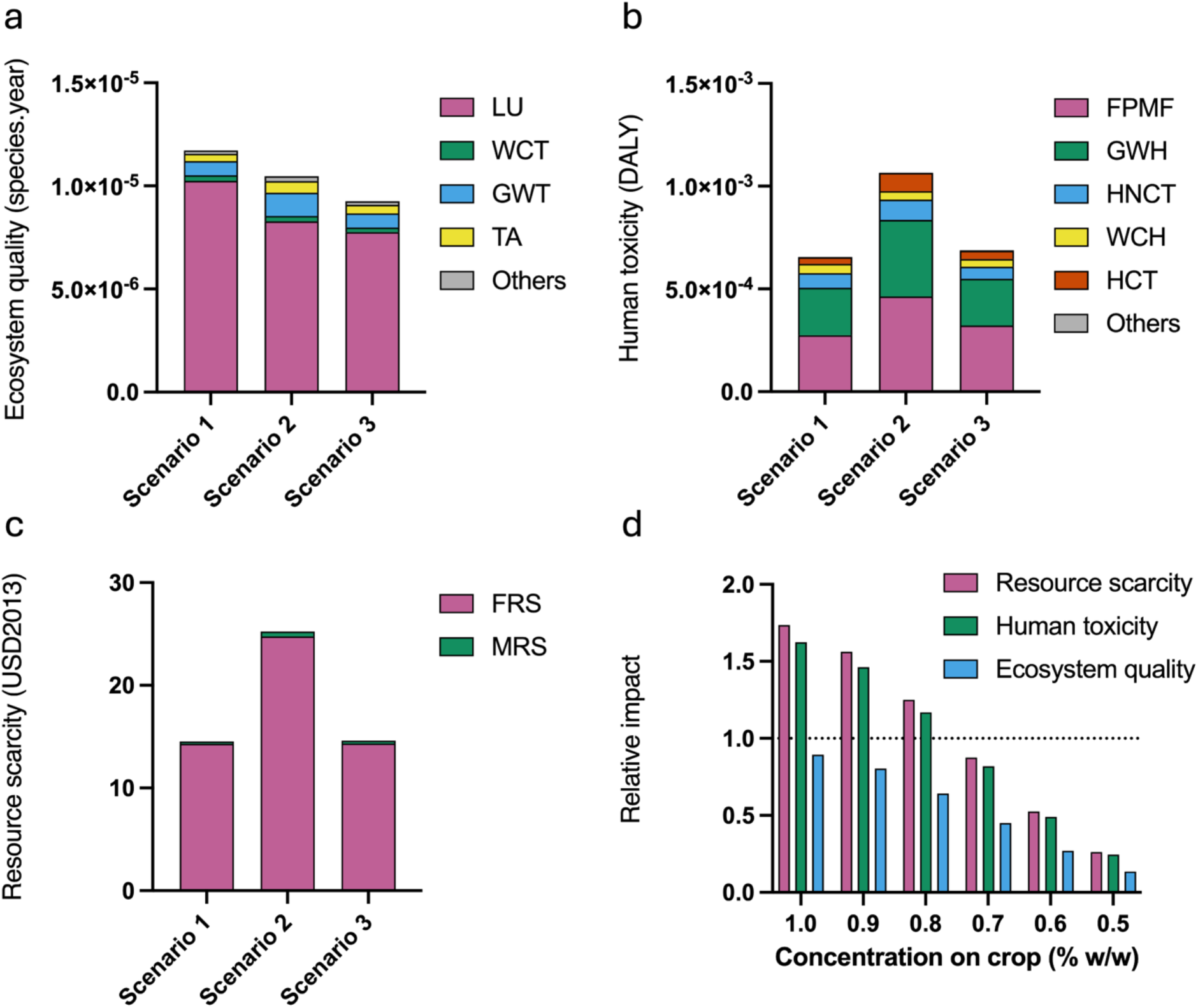
Impact of the three scenarios on (a) ecosystem quality, (b) human toxicity, and (c) resource scarcity. (d) A sensitivity analysis of scenario 2 using scenario 1 as the baseline (dashed line) on the three endpoint categories. LU, land use; WCT, water consumption terrestrial ecosystems; GWT, global warming terrestrial ecosystems; TA, terrestrial acidification; FPMF, fine particle matter formation; GWH, global warming human health; HNCT, human non carcinogenic toxicity; WCH, water consumption human health; HCT, human carcinogenic toxicity; FRS, fossil resource scarcity; MRS, mineral resource scarcity. Others comprise the impact categories not represented in the legend (Supplementary Table 8).

Due to limited resources, we have not been able to conduct a bird repellence dose-response curve of MANT-*N*-glucose in this study. Since we saw 100 % repellence at the chosen dose (1.0 % w/w), it is possible that lower amounts of MANT-*N*-glucose can be used while preserving efficiency. A sensitivity analysis of the end-point impacts showed that for chemoenzymatic MANT-*N*-glucose to be an eco-friendlier choice, it should be applied at a concentration equal to or lower than 0.7 % (w/w) (Figure 5d).

The significant impacts of chemoenzymatic MANT-*N*-glucose can be attributed to UDP-glucose and DMSO (Supplementary Figure 16). Although the UDP-glucose recycling system based on SuSy is a well-established method to eliminate the need for stoichiometric UDP-glucose,^[59]^ the widely used *Gm*SuSy was inhibited by the substrate for the glycosylation reaction, MANT, or by the DMSO used to solubilise MANT (Figure 3a). Microbial production from a sugar feedstock is another approach that would address the challenges associated to the use of UDP-glucose and DMSO. Consequently, we modelled a microbial cell factory system (scenario 3) based on a published *C. glutamicum* strain producing MANT at a titre of 5.7 g/L.^[20]^ Based on our antimicrobial activity data (Figure 4), we assumed low toxicity of MANT-*N*-glucose on *C. glutamicum*, allowing for an identical titre to the published one, without using the two-phase extraction system used in the publication (Supplementary Table 4).^[20]^ With this approach, microbial MANT-*N*-glucose demonstrated a reduction in seven impact categories when compared to chemically synthesised MANT (Supplementary Table 7), with significant improvements in land use and marine eutrophication (26 % decrease in both). The primary contributor in this scenario is the glucose feedstock, accounting for approximately 50 % of the total impact of the main toxicity indicators, except for human non-carcinogenic toxicity, where it contributed 10 % (Supplementary Figure 16). At the endpoint level, microbial MANT-*N*-glucose outperformed chemoenzymatic MANT-*N*-glucose in all three categories (Figure 5). In contrast, it performed similarly to chemically synthesised MANT in both human toxicity and resource scarcity while showing a 20 % reduction in the ecosystem quality impact (Figure 5).

Overall, the environmental implications of producing and using chemoenzymatic MANT-*N*-glucose have been identified and compared to the existing product, chemically synthesised MANT, as well as to a hypothetical microbially synthesised MANT-*N*-glucose. The use of stochiometric amounts of UDP-glucose is a key environmental driver, which could be addressed by the discovery or engineering of a more robust SuSy that would allow efficient UDP-glucose recycling even at high MANT titres. A limitation of our models is the lack of detailed purification processes, which often represent the most significant impact on the environmental and economic performance of a bioprocess.^[60]^ In terms of the microbially produced MANT-*N*-glucose, the 20% lower biodiversity impact and the similar human toxicity and resource scarcity impacts relative to chemical MANT make this strategy a promising replacement for MANT as a bird deterrent, especially if a lower dose can be applied due to the higher bird repellence of the glucoside.

## Conclusion

In this study, we report the first UGTs active on AEANTs. Among the three candidate UGTs with the broadest AEANT specificity, UGT72B68 exhibited the highest catalytic efficiency with MANT, leading to its selection for further experiments. Rational engineering of its active site produced a variant, F145M, with improved activity with all AEANTs besides MANT. Furthermore, UGT72B68 displayed excellent chemo tolerance, which enabled the multigram-scale production of pure MANT-*N*-glucose. The purified product displayed decreased antimicrobial effects on *E. coli* or *P. putida* and exhibited efficient bird repellence. A preliminary LCA highlighted the limitations of using MANT-*N*-glucose both as a bird repellent and as a fermentation target, providing insights into its environmental trade-offs and indicating microbial production as a way forward. Overall, this study showcased a complete pathway from enzyme discovery to practical application, providing a foundation for future research and development in the field of biocatalysis. Additionally, the potential of glycosylating bioactive compounds was demonstrated to go beyond enhancing their physicochemical properties but also as a tool to enable the synthesis of novel compounds with improved biological activity.

## Supporting Information

The authors have cited additional references within the Supporting Information. ^[61–67]^

## Data Availability Statement

The data that support the findings of this study are available in the supplementary material of this article.

## Acknowledgements

We thank The Novo Nordisk Foundation for supporting this work through grants NNF20CC0035580 and NNF22SA0078231. We thank Daniela Rago and Onur Kırtel from The Novo Nordisk Foundation Center for Biosustainability for their contributions to the LC-MS measurements and the LCA analysis.

## Entry for the Table of Contents

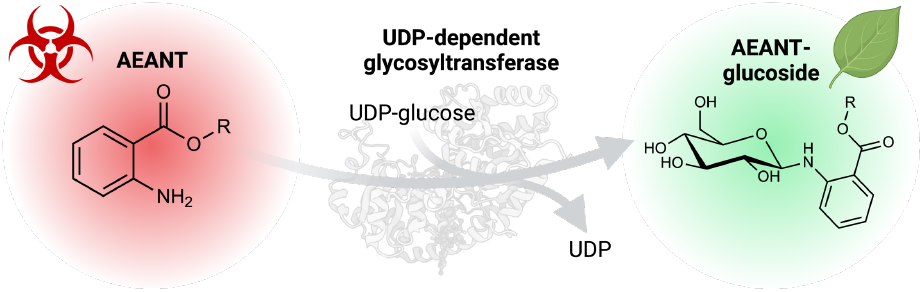

We report the discovery of UDP-dependent glycosyltransferases (UGTs) capable of glycosylating alkyl-ester derivatives of anthranilic acid (AEANTs), thereby mitigating their undesirable physicochemical properties and antimicrobial activity. We scaled-up enzymatic synthesis of methyl anthranilate *N*-glucose and evaluated its antimicrobial and bird-repellent effects, demonstrating the functional benefits of AEANT glycosylation.

## Notes

Supporting information for this article is given via a link at the end of the document.

### Competing Interest Statement

The authors have declared no competing interest.

