## Supplementary Information for "Enzymatic Glycosylation of Anthranilates for Enhanced Functionality"

The supplementary information includes:

**Materials and Methods**

**Supplementary Tables**

Supplementary Table 1. Bird repellence study timeline
Supplementary Table 2. Mass and energy balance, Scenario 1
Supplementary Table 3. Mass and energy balance, Scenario 2
Supplementary Table 4. Mass and energy balance, Scenario 3
Supplementary Table 5. Assumptions used in the life cycle inventory
Supplementary Table 6. UGT AEANT activity screening
Supplementary Table 7. Impact on midpoint categories
Supplementary Table 8. Impact on endpoint categories

**Supplementary Figures**

Supplementary Figure 1. System boundaries and flow diagram of the preliminary LCA analysis
Supplementary Figure 2. Optimal UGT reaction pH
Supplementary Figure 3. Optimal UGT reaction temperature.
Supplementary Figure 4. Kinetic characterisation, MANT
Supplementary Figure 5. Kinetic characterisation, EANT, BANT, and IBANT
Supplementary Figure 6. Molecular docking poses
Supplementary Figure 7. Active site residues, UGT72B68 and UGT71C1
Supplementary Figure 8. Chai-1 predicted binding poses
Supplementary Figure 9. Biocatalytic MANT-*N*-glucose chromatogram
Supplementary Figure 10. MANT-*N*-glucose H-NMR
Supplementary Figure 10. MANT-*N*-glucose C-NMR
Supplementary Figure 12. MANT-*N*-glucose HMBC
Supplementary Figure 13. MANT-*N*-glucose chemical degradation
Supplementary Figure 14. MANT-*N*-glucose chemical degradation products
Supplementary Figure 15. LC-MS validation of degradation product
Supplementary Figure 16. Contributors of the main impact categories

### Materials and Methods

#### Materials

Buffers, chemicals, and reagents were purchased from commercial vendors. Point mutation variants of UGT72B68 were synthesised by Biomatik (USA). UDP-glucose was purchased from Ambeed, Inc. (USA). MANT-*N*-glucose was not commercially available during the discovery, characterisation, and engineering phases.

#### Methods

**Cloning, expression, and purification of recombinant UGTs**

The full-length coding sequence of the UGTs, along with variants, were synthesised and cloned into the pET28a(+) vector by Genscript (USA) or Biomatik (USA), using the NcoI and XhoI restriction sites. The protein-coding sequences included an N-terminal 6xHis-tag and a TEV-cleavage site. Plasmids were transformed into *E. coli* BL21 Star™ (DE3) cells (Thermo Fisher Scientific). Precultures were prepared by inoculation from glycerol stocks in 15 mL 2xYT medium supplemented with 50 μg $mL^{-1}$ kanamycin and grown overnight at 37 °C, 250 rpm. Precultures were then diluted to $OD_{600}$ = 0.05 in 2xYT supplemented with 50 μg $mL^{-1}$ kanamycin in a baffled flask to a final volume of 1000 mL. The cultures were incubated at 37 °C, 180 rpm, until the $OD_{600}$ reached 0.6-0.8 upon which it was induced with 0.2 mM IPTG and subsequently incubated overnight at 20 °C, 180 rpm. The cells were harvested by centrifugation at 4500 g, 30 min at 4 °C. The cell pellet was washed with phosphate-buffered saline (PBS) pH 7.4 once followed by resuspension in an appropriate amount (10 mL pr. g of the pellet) of lysis buffer (50 mM phosphate, 300 mM NaCl, 20 mM imidazole, pH 7.5, supplemented with 10 μg/mL DNase, 0.1 mg/mL lysozyme, 0.05 % (v/v) Triton X-100). Cell lysis was performed by sonication on a Sonics Vibra Cell (USA) at 85 % amplitude with 15 seconds on/30 seconds off intervals for a total of 4 minutes of sonication time. The soluble cell lysate was recovered by centrifugation at 14,500 × g at 4 °C for 45 min. The supernatant containing the recombinant proteins was filtered with a 0.45 μm filter and purified by nickel-affinity chromatography (prepacked HisTrap™ FF columns, GE Healthcare) on an ÄKTA pure (GE Healthcare). Before sample loading, the column was washed and equilibrated with buffer A (50 mM phosphate, 300 mM NaCl, pH 7.5, with 20 mM imidazole). Following protein binding, the column was washed with buffer A and subsequently eluted using a gradient elution with buffer A and buffer B (50 mM phosphate, 300 mM NaCl, pH 7.5, with 500 mM imidazole). The elution peak fractions were pooled together and concentrated using centrifugal filters to a volume of 2.5 mL. The buffer was exchanged for the storage buffer (50 mM phosphate, 150 mM NaCl, pH 7.5) via PD10 desalting columns packed with Sephadex G-25 resin (Cytiva). The total concentration of the purified proteins was measured by spectrophotometric measurements at 280 nm using a NanoDrop 2000 (Thermo Fisher Scientific) and adjusted using the extinction coefficient of the respective proteins. The sample purity was assessed using sodium dodecyl sulfate-polyacrylamide gel electrophoresis (SDS-PAGE).

**HPLC analysis for enzyme assays and degradation studies**

Product formation was followed by HPLC, using an Ultimate 3000 Series apparatus (Thermo Fisher) employing a Kinetex C18 analytical column (dimensions 4.6 x 100 mm; pore size 100 Å; particle size 2.6 µm) (Phenomenex). 0.1 % formic acid in water (A) and acetonitrile (B) were used as mobile phases employing a gradient elution*:* 0.00–0.50 min, 2% B;0.50-0.51 min, linear ramp to 35 % B; 0.51–1.90 min, 35 % B; 1.90-2.50 min, linear ramp to 100 % B; 2.50–4.20min, 100 % B; 4.20-4.21, linear ramp to 2 % B; 4.21-5.00 min, 2 % B. The flow rate was set to 1 mL/min. The analytes were detected at 330 nm. The HPLC data was monitored and quantified via the Chromeleon software (Thermo Fisher Scientific). The conversion was determined for the enzyme discovery assays by analysing the ratio between the substrate in the reaction containing enzyme and the control reaction without enzyme. For the remaining assays, unless stated otherwise, the formation of the product was determined by analysing the ratio between the product peak and acceptor peak on the HPLC chromatograms, assuming the absorbance at 330 nm is equal.

**HPLC analysis for the multigram-scale production of MANT-*N*-glucose**

Fractions and final products were analysed via HPLC, using an Ultimate 3000 Series apparatus (Thermo Fisher) employing a Zorbax Eclipse Plus C18 analytical column (dimensions 4.6 x 100 mm; pore size 95 Å; particle size 3.5 µm). 0.1 % formic acid and 10 mM ammonium formate in water (A) and acetonitrile (B) were used as mobile phases employing a gradient elution: 0.00-2.00 min, 10 % B; 2.00-10.00 min, linear ramp to 70 % B; 10.00-12.00 min, 70 % B; 12.00-13.00 min, linear ramp to 10 % B; 13.00-15.00 min, 10 % B. The flow rate was set to 1 mL/min. The analytes were detected at 260 nm and 330 nm. The HPLC data was monitored and quantified via the Chromeleon software (Thermo Fisher Scientific). An authentic MANT-*N*-glucose standard (Biosynth) was used for validation.

**LC-MS/MS analysis**

The LC-MS/MS analysis was performed as previously described.[1] Briefly, the LC-MS/MS analysis was performed on a Vanquish Duo UHPLC binary system (Thermo Fisher Scientific) coupled to an IDX-Orbitrap mass spectrometer (Thermo Fisher Scientific). The chromatographic separation was achieved using a Waters ACQUITY BEH C18 (dimensions 2.1 x 100 mm; pore size 130 Å; particle size 1.7 µm) equipped with an ACQUITY BEH C18 guard column kept at 40 °C. 0.1 % formic acid in water (A) and 0.1 % formic acid in acetonitrile (B) were used as mobile phases employing a gradient elution: 0.00-0.80 min, 2 % B; 0.80-3.30 min, linear ramp to 5 % B; 3.30-10.00 min, linear ramp to 100 % B; 10.00-11.00 min, 100 % B. The re-equilibration time was 2.7 min. The flow rate was set to 0.35 mL/min. The MS measurement was done in positive—heated electrospray ionization (HESI) mode with a voltage of 3,500 V acquiring full MS/MS spectra (data-dependent acquisition-driven MS/MS) in the mass range of 70–1,000 Da.

**Sequence selection and initial activity assays**

The initial activity screening utilised an in-house library[2] of 45 sequences randomly selected from plant species with known soluble expression. From these 45 sequences, 19 were randomly chosen for the MANT assay, and 28 were selected for the EANT assay.

The MANT and EANT activity assays were performed in 50 mM 4-(2-hydroxyethyl)-1-piperazineethanesulfonic acid (HEPES) pH 7.0 and consisted of a nonspecific amount of the UGT, 1 mM UDP-glucose, and 0.1 mM of the acceptor from a 50 mM dimethyl sulfoxide (DMSO) stock. After an overnight reaction at room temperature, the mixtures were centrifuged at 17,000 × g for 5 min, and the supernatant was analysed via HPLC.

UGTs exhibiting > 50 % conversion with MANT and/or EANT were further tested with BANT. This assay was also performed in 50 mM HEPES pH 7.0 and 1 mM UDP-glucose but was assayed with 0.5 mM BANT from a 50 mM DMSO stock and 2.5 µM enzyme. The three UGTs showing the highest activity with MANT, EANT, and BANT were subsequently assayed with IBANT and CANT, following the same protocol used for BANT. A reaction mixture without UGT was used as a control in all assays.

**Biochemical characterisation with MANT**

*pH optimum*

Reactions were performed at a pH range to find the optimum pH of UGT72B68, UGT72B19, and UGT71C1 using MANT as the acceptor substrate. The following buffering agents were utilised for the respective pH ranges: pH 4-7, 50 mM citrate-phosphate; pH 7-8, 50 mM phosphate; pH 8-9, 50 mM Trizma; pH 9-11, 50 mM glycine. Each reaction contained 0.40 mM UDP-glucose, 0.11 mM MANT from a 50 mM DMSO stock, and UGT (0.7 µM for UGT72B68 and 1.1 µM for UGT72B19 and UGT71C1). The reaction was incubated at room temperature and quenched at 5 time points (2 min, 5 min, 10 min, 15 min, and 30 min) by transferring 50 µL of the reaction mixture to 70 µL of cold methanol. The reaction mixtures were analysed via HPLC. The initial rates were obtained and plotted as means ± standard deviations of duplicate experiments.

*Temperature optimum*

Reactions were performed at a temperature range to find the optimum temperature of UGT72B68, UGT72B19, and UGT71C1 using MANT as the acceptor substrate. Each reaction contained 0.40 mM UDP-glucose, 0.11 mM MANT from a 50 mM DMSO stock, and UGT (0.7 µM for UGT72B68 and 1.1 µM for UGT72B19 and UGT71C1).

50 mM phosphate buffer at the previously determined optimal pH was used and the reaction was carried out at temperatures ranging from 30 °C to 54 °C for UGT72B68 and UGT72B19 and 15 °C to 37 °C for UGT71C1 on a thermocycler. The reaction was stopped at four different time points (5 min, 10 min, 15 min, and 30 min) by thermal denaturation at 95 °C for 20 seconds. The reaction mixtures were analysed via HPLC. The initial rates were obtained and plotted as means ± standard deviations of duplicate experiments.

**Kinetic characterisation of UGT72B68, UGT72B19, and UGT71C1 with MANT**

The kinetic properties of UGT72B68, UGT72B19, and UGT71C1 with MANT were determined in 50 mM phosphate buffer at the apparent pH and temperature optima for the respective UGTs. Reaction mixtures consisted of 0.8 mM UDP-glucose, an appropriate UGT concentration (0.7 μM for UGT72B68, 1.1 μM for UGT72B19, and 0.5 μM for UGT71C1), and a MANT concentration ranging from 1000 μM to 11.56 μM from a 50 mM stock in DMSO. Reactions were performed on a thermocycler. The reaction was stopped at four different time points (2 min, 5 min, 10 min, and 15 min) by thermal denaturation at 95 °C for 20 seconds. The reaction mixtures were analysed via HPLC. The initial rates were obtained and plotted as means ± standard deviations of duplicate experiments. Michaelis-Menten plots were generated and analysed in Prism v10.4.0.

**Kinetic characterisation of UGT72B68 with EANT, BANT, and IBANT**

The kinetic properties of UGT72B68 with EANT, BANT, and IBANT were determined using the apparent pH and temperature optima achieved with MANT (50 mM phosphate buffer pH 8.0 at 35 °C). For EANT, reaction mixtures consisted of 0.40 mM UDP-glucose, 0.40 μM UGT72B68, and an EANT concentration ranging from 2000 μM to 23.12 μM from a 50 mM stock in DMSO. The reaction was stopped at four different time points (2 min, 5 min, 10 min, and 15 min). For BANT and IBANT, reaction mixtures consisted of 0.40 mM UDP-glucose, 0.25 μM UGT72B68, and a BANT/IBANT concentration ranging from 1000 μM to 11.56 μM from a 10 mM stock in DMSO (lower water solubility compared to MANT). The reaction was stopped at four different time points (1 min, 2 min, 3 min, and 5 min). Reactions were stopped by thermal denaturation at 95 °C for 20 seconds. The reaction mixtures were analysed via HPLC. The initial rates were obtained and plotted as means ± standard deviations of duplicate experiments. Michaelis-Menten plots were generated and analysed in Prism v10.4.0.

**Molecular docking and enzyme-substrate complex modelling**

MANT, EANT, BANT, IBANT, and CANT were docked into an AlphaFold[3] model of UGT72B68 using AutoDock Vina v1.1.2,[4] utilising standard settings. To help define the grid space, a binary complex of UGT72B68 and UDP-glucose was obtained by structurally aligning the UGT72B68 AlphaFold model on the crystal structure of

*Pt*UGT1 from *Polygonum tinctorium*, which has a bound UDP-glucose molecule in its active site (PDB: 6SU6).[5] The grid space was placed near the glucose moiety of UDP-glucose, and docked poses were inspected in PyMOL (v2.5.2). Poses where the aniline group of AEANT was not oriented toward UDP-glucose, particularly its anomeric carbon, were excluded from consideration. The docked poses were validated using Chai-1 with MSAs enabled.[6] The sequence of UGT72B68 was inputted along with the smiles code of UDP-glucose and one of the six respective AEANTs. Complexes where the aniline group of AEANT was not oriented toward UDP-glucose, particularly its anomeric carbon, were excluded from consideration.

**Relative conversion yield of UGT72B68 variants**
Reaction mixtures contained 2.0 mM UDP-glucose, 1.0 mM acceptor substrate from a 10 mM stock in DMSO, and 2.5 µM of the UGT variant in 50 mM phosphate buffer pH 8.0 at 35 °C for 15 minutes. Reactions were stopped by thermal denaturation at 95 °C for 20 seconds. The reaction mixtures were analysed via HPLC as means ± standard deviations of two independent experiments. The activity was quantified via the product peak area and subsequently normalised to the variant with the largest peak area for each substrate, respectively.

**Chemotolerance of *Gm*SuSy and UGT72B68 towards MANT**
The chemo tolerance of *Gm*SuSy was tested by reacting 0.25-1.0 µM *Gm*SuSy with 1.0 mM UDP, 100 mM sucrose, 0.75 µM UGT72B68, and 0.12-10 mM MANT from a 50 mM DMSO stock in 50 mM phosphate buffer pH 8.0 at 35 °C for 2, 5, 10, and 15 minutes. Reactions were stopped by thermal denaturation at 95 °C for 20 seconds. The initial rates were obtained and plotted as a single replicate experiment.

The chemo tolerance of UGT72B68 was first tested by reacting 0.5, 1.0, and 1.5 µM UGT72B68 with 1.0 mM UDP-glucose and 0.12-10 mM from a 50 mM DMSO stock in 50 mM phosphate buffer pH 8.0 at 35 °C for 2, 5, 10, and 15 minutes. The experiment was performed as a single replicate. Then, the chemo tolerance of UGT72B68 was further probed by reacting 0.75 µM UGT72B68 with 1.0 mM UDP-glucose and 0.23-20.0 mM from a 100 mM DMSO stock in 50 mM phosphate buffer pH 8.0 at 35 °C for 2, 5, 10, and 15 minutes. The initial rates were obtained and plotted as means ± standard deviations of duplicate experiments. All reaction mixtures were analysed via HPLC.

**Production and purification of multigram-scale MANT-glucose**
After establishing the tolerance of UGT72B68 towards MANT, all UGT72B68 produced from 22 L of culture was utilised in a 2 L reaction volume. This resulted in a mixture of 20 mM MANT (from a 200 mM DMSO stock), 30 mM UDP-glucose, and 2.3 µM UGT72B68. The reaction was performed in a 2 L flask and incubated at 35 °C at 90

RPM. The reaction mixture was incubated for 20 hours, after which a sample was taken and analysed via HPLC.

The reaction mixture was filtered through a 0.22 μm PES sterile filter. The separation of the product from the substrate, UDP, UDP-glucose, and the remaining enzyme was achieved on a Flash Buchi C-850 system using an FP ECOFLEX C18 40g column. Approximately 300 mL of the reaction mixture was injected into the column during each run. Two mobile phases (A: MQ water and B: 90:10 MeOH:MQ water) were utilised in a gradient elution: 0.00–2.90 min, 10 % B; 2.90–28.60 min, linear ramp 10–90 % B; 28.60–34.30 min, 90 % B; 34.30–35.70 min, linear ramp 90–10 % B; 35.70–40.00 min, 10 % B. The flow rate was set to 45 mL/min. MANT and MANT-N-glucose were detected via UV at 330 nm, while UDP and UDP-glucose were detected at 265 nm. Isolated fractions were combined and analysed via HPLC. The collected fractions were evaporated at 50 °C using a Buchi Rotavapor R-300 to separate methanol and water from the product. Methanol evaporated at 150 mbar, and water evaporated at 50 mbar. After reaching the critical solubility of the product in water (after evaporating approximately 1 L of methanol and water), the product precipitated, and the evaporation was stopped. The condensate, which contained some product, was reevaporated to maximise the product recovery. The precipitate was filtered and freeze-dried for 48 hours. The purity was measured via HPLC, and the product was validated via NMR using DMSO-d6.

**Bird repellent effects of MANT-*N*-glucose on red-winged blackbirds**

One feeding experiment was conducted in February 2024 at the United States Department of Agriculture, National Wildlife Research Center's (NWRC) outdoor animal research facility in Fort Collins, Colorado (USA). 11 male red-winged blackbirds (*Agelaius phoeniceus*) were live-captured for the experiment. The capture, care, and use of all birds associated with the feeding experiment were approved by the NWRC Animal Care and Use Committee (NWRC Study Protocol QA-3596; A.K. Kohler- Study Director).

Blackbirds were maintained in a 4.9 × 2.4 × 2.4 m cage within a wire mesh-sided building for at least two weeks before the experiment (i.e., quarantine, holding; Supplementary Table 1). Free access to grit (sand) and a maintenance diet was provided to all birds during quarantine and holding. The maintenance diet consisted of two parts millet and one part cracked corn, milo, and safflower. The experiment was conducted in visually isolated individual cages (0.9 × 1.8 × 0.9 m) within a wire mesh-sided building. Water was provided ad libitum to all birds throughout the experiment (quarantine, holding, preference experiment).

The seed treatments offered during the experiment were formulated by applying aqueous suspensions (60 mL/kg) to whole oilseed sunflower (Northern Colorado

Feeders Supply, Fort Collins, CO, USA) using a rotating mixer and household spray equipment. All blackbirds were offered untreated sunflower seeds ad libitum in two food bowls for five days of acclimation in individual cages (Supplementary Table 1). Each blackbird was subsequently offered one bowl of untreated sunflower and one bowl of sunflower treated with 1 % MANT-*N*-glucose (targeted concentration, w/w) at 08:00 h during the test (i.e., Monday; Supplementary Table 1). Daily sunflower consumption was measured during the one-day preference experiment. Unconsumed sunflower seeds (remaining in each food bowl) and spillage were collected (at 08:00 h on the day after the test; Tuesday) and weighed (± 0.1 g). Weight change (*e.g.*, desiccation) of sunflower seeds was measured daily by weighing seeds offered within a vacant cage throughout the preference experiment.

The dependent measure for the preference experiment was the average (i.e., daily) test consumption of treated and untreated sunflower seeds. Descriptive statistics (± SEM) and a paired t-test were used to summarise and analyse the consumption of treated and untreated seeds throughout the experiment.

**Antimicrobial activity of MANT and MANT-*N*-glucose on *E. coli* and *P. putida***
Precultures of *E. coli* BL21 Star™ (DE3) cells and *P. putida KT2440* were grown in Luria Bertani (LB) media overnight at 37 °C and 30 °C, respectively, with agitation (200 rpm). Overnight cultures were diluted to $OD_{600}$ = 0.10 in M9 minimal media with or without DMSO/MANT/MANT-*N*-glucose. MANT and MANT-*N*-glucose were added at concentrations 1.5-7.5 mM from a 150 mM DMSO stock. DMSO was added to all samples to match the DMSO concentration of the sample containing 7.5 mM MANT(-Glc) (5% v/v). A control without substrate and 5 % (v/v) DMSO and a control without substrate and 0 % (v/v) DMSO were also prepared. 200 μL cultures were incubated in 96-well plates sealed with a Breathe-Easy® (Sigma-Aldrich, USA) sealing membrane at 37 °C and 30 °C for *E. coli* and *P. putida*, respectively, with agitation (567 cpm) in an Epoch2 microplate spectrophotometer (BioTek, Agilent, USA). Bacterial growth was monitored every 10 minutes for 24 hours by measuring the $OD_{600}$. The data was analysed as means ± standard deviations of duplicate experiments.

Samples with MANT-*N*-glucose concentrations ranging from 10-50 mM were also prepared using an 80 mM stock of MANT-*N*-glucose in water. Here, no DMSO was added. Otherwise, all other conditions and analysis methods were kept as described above.

**The chemical degradation of MANT-*N*-glucose**
The chemical degradation of MANT-*N*-glucose was investigated in acidic and alkaline conditions. 5 mM of MANT-*N*-glucose from a 50 mM stock in water was added to aqueous solutions containing 1 mM, 10 mM, or 100 mM HCl or NaOH, respectively. A control sample in water, without acid or base, was monitored simultaneously. Samples

were collected at 0, 1, 2, 3, 4, and 5 hours, as well as after an overnight incubation. Samples were diluted 20 times in 50 mM phosphate buffer pH 7.0 to neutralise the acidity/alkalinity and subsequently stored at -18°C until further analysis. The reaction mixtures were analysed via HPLC, and the MANT-*N*-glucose abundance was evaluated relatively by comparing the peak areas of MANT/MANT-*N*-glucose/ANT/ANT-*N*-glucose. The data was analysed as means ± standard deviations of duplicate experiments.

**Preliminary life-cycle assessment**

*Functional unit and system boundaries*

The functional unit was defined as the production of 100 kg of sunflower seeds using a treatment with MANT or MANT-*N*-glucose at a 1 % w/w concentration and correcting for the difference in bird repellence by MANT (70 %)[7] and MANT-*N*-glucose (100 %). The system boundaries were established according to the operational units of three scenarios, each employing a distinct production of bird repellent (Supplementary Figure 1): chemical synthesis of MANT (scenario 1), chemoenzymatic synthesis of MANT-*N*-glucose (scenario 2), and microbial synthesis of MANT-*N*-glucose (scenario 3). A cradle-to-gate system was modelled by considering the mass balances for each operational unit.

*Life-cycle inventory and impact assessment*

The life cycle inventory (LCI) of each operational unit was based on literature data, including research articles and patents referenced throughout the paper, except for MANT glycosylation, which was investigated in this study. The modelled chemical synthesis of MANT consisted of incubating ANT with an excess of methanol using an acid catalyst (cationic resin) at 120 °C for 6 hours (Supplementary Table 2).[8] Chemoenzymatic synthesis of MANT-*N*-glucose was modelled with 30 mM UDP-glucose, 20 mM MANT, and 2.30 µM UGT72B68 (Supplementary Table 3). These amounts matched those used for the production and purification of MANT-*N*-glucose described in this study. UDP-glucose synthesis was modelled based on the whole-cell conversion of UDP and sucrose by *E. coli* overexpressing SuSy from *Acidithiobacillus caldus*.[9] Meanwhile, UDP synthesis was modelled based on the whole-cell biocatalytic conversion of uridine using *E. coli* and *Saccharomyces cerevisiae*.[10] Microbial MANT-*N*-glucose synthesis was modelled using the fermentation conditions of MANT production from *Corynebacterium glutamicum* (Supplementary Table 4).[11] Due to the difference in the bird repellence of MANT and MANT-*N*-glucose, the use of chemically synthesised MANT resulted in a production loss of 40 kg. As this is a preliminary LCA based on laboratory data, utilities, and purification processes for MANT and MANT-*N*-glucose were not considered. Assumptions were made due to the limited availability of data for biological processes in the database (Supplementary Table 5). The life cycle impact assessment (LCIA) was calculated using openLCA 2.1 with the ecoinvent v3.8 database as the source of background systems and ReCiPe

2016 Midpoint (H) and Endpoint (H) as the impact assessment method with normalisation against the World 2010 (H) database.

### Supplementary Tables

| Experimental phase | Timeline |
|---|---|
| Quarantine/Holding | 2+ weeks |
| Acclimation | Wednesday-Sunday (5 days) |
| Test | Monday (1 day) |

**Supplementary Table 1. Bird repellence study timeline.** Experimental timeline describing the days or weeks for each experimental phase of the bird repellence study.

| Flows | Unit | Quantity | Provider in openLCA | Allocation |
|---|---|---|---|---|
| **Inputs** | | | | |
| Sunflower seeds | kg | 140.000 | Market for sunflower seed | GLO |
| Anthranilic acid | kg | 0.907 | Market for anthranilic acid | GLO |
| Methanol | kg | 7.187 | Market for methanol | GLO |
| Acid catalyst | kg | 0.245 | Market for cationic resin | RER |
| **Outputs** | | | | |
| Hazardous waste | kg | 7.339 | Waste incineration with energy recovery | EWS |
| Sunflower seeds loss | kg | 40.000 | | |
| Sunflower seeds | kg | 100.000 | | |

**Supplementary Table 2. Mass and energy balance, Scenario 1.** Mass and energy balance for the production of 100 kg of sunflower seeds using chemically synthesised MANT (Scenario 1).[8] EWS, Europe without Switzerland; RER, region of Europe; GLO, global; DK, Denmark.

| Flows | Unit | Quantity | Provider in openLCA | Allocation |
|---|---|---|---|---|
| **Inputs** | | | | |
| Sunflower seeds | kg | 100.000 | Market for sunflower seed | GLO |
| Enzyme | kg | 0.021 | Market for enzymes | GLO |
| Buffer | kg | 11.579 | Market for sodium phosphate | RER |
| DMSO | kg | 19.506 | Market for dimethyl sulfoxide | GLO |
| Water | kg | 351.955 | Market for water, deionized | EWS |
| Glucose | kg | 19.810 | Market for glucose | GLO |
| Yeast extract | kg | 0.280 | Market for protein feed, 100 % crude | GLO |
| Tryptone | kg | 0.186 | Market for protein feed, 100 % crude | GLO |

| | | | | |
|---|---|---|---|---|
| $MgSO_4$ | kg | 0.282 | Market for magnesium sulphate | GLO |
| $FeSO_4$ | kg | 0.0003 | Market for iron sulphate | GLO |
| $MnSO_4$ | kg | 0.0003 | Market for manganese sulphate | GLO |
| Brewer's yeast | kg | 39.172 | Market for fodder yeast | GLO |
| NaCl | kg | 0.042 | Market for sodium chloride | GLO |
| $NH_4Cl$ | kg | 0.003 | Market for ammonium chloride | GLO |
| Sucrose | kg | 16.269 | Market for sugar, from sugar beet | GLO |
| MANT | kg | 0.536 | | |
| **Outputs** | | | | |
| Biowaste | kg | 43.312 | Municipal incineration | GLO |
| Wastewater | L | 416.329 | Wastewater, average, capacity $10^9$ L | EWS |
| Sunflower seeds | kg | 100.000 | | |

**Supplementary Table 3. Mass and energy balance, Scenario 2.** Mass and energy balance for the production of 100 kg of sunflower seeds using chemoenzymatically produced MANT-*N*-glucose (Scenario 2).[9,10,12] EWS, Europe without Switzerland; RER, region of Europe; GLO, global; DK, Denmark.

| Flows | Unit | Quantity | Input in openLCA | Allocation |
|---|---|---|---|---|
| **Inputs** | | | | |
| Sunflower seeds | kg | 100.000 | Market for sunflower seed | GLO |
| $(NH_4)_2SO_4$ | kg | 5.323 | Market for ammonium sulphate | RER |
| Urea | kg | 0.271 | Market for urea | RER |
| Buffer | kg | 0.388 | Market for sodium phosphate | RER |
| $MgSO_4$ | kg | 0.032 | Market for magnesium sulphate | GLO |
| $CaCl_2$ | kg | 0.002 | Market for calcium chloride | RER |
| $FeSO_4$ | kg | 0.002 | Market for iron sulphate | GLO |
| $MnSO_4$ | kg | 0.002 | Market for manganese sulphate | GLO |
| $ZnSO_4$ | kg | 0.0002 | Market for zinc sulfide | GLO |
| $CuSO_4$ | kg | 0.00006 | Market for copper sulphate | GLO |
| Glucose | kg | 65.525 | Market for glucose | GLO |
| Yeast extract | kg | 2.662 | Market for protein feed, 100 % crude | GLO |
| Water | kg | 266.550 | Market for water, deionized | EWS |

| **Outputs** | | | | |
|---|---|---|---|---|
| Biowaste | kg | 1.333 | Municipal incineration | GLO |
| Wastewater | L | 338.472 | Wastewater, average, capacity $10^9$ L | EWS |
| Sunflower seeds | kg | 100.000 | | |

**Supplementary Table 4. Mass and energy balance, Scenario 3.** Mass and energy balance for the production of 100 kg of sunflower seeds using microbial MANT-*N*-glucose based on the 2-phase system titer (Scenario 3).[11] EWS, Europe without Switzerland; RER, region of Europe; GLO, global; DK, Denmark.

| **Input** | **Process** | **Included as** |
|---|---|---|
| Yeast extract | Uridine<br>UDP-glucose<br>Microbial MANT-*N*-glucose | Protein feed, 100 % crude |
| Tryptone | Uridine<br>UDP-glucose | Protein feed, 100 % crude |
| $KH_2PO_4$ | Uridine<br>UDP-glucose<br>Microbial MANT-*N*-glucose | Sodium phosphate |
| $K_2HPO_4$ | UDP-glucose<br>Microbial MANT-*N*-glucose | Sodium phosphate |
| Brewer’s yeast | UDP | Fodder yeast |
| $ZnSO_4$ | Microbial MANT-*N*-glucose | Zinc sulfide |

**Supplementary Table 5. Assumptions used in the life cycle inventory.** These inputs were not found in the database, so other inputs with similar functions were chosen, assuming that a similar impact could be obtained.[13] Vitamins B1, B3, B5, B7, and B12 together with $NiCl_2$, tryptophane, polypropylene glycol, protocatechuic acid, antibiotics, and IPTG were not included in the LCI since they were not found in the database, and they account for < 0.1 % of the total input of the process.

| **UGT isoform** | **MANT** | **EANT** | **BANT** | **IBANT** | **CANT** |
|---|---|---|---|---|---|
| **UGT706F8** | 46.4 | 31.1 | NT | NT | NT |
| **UGT706H1** | NT | 16.6 | NT | NT | NT |
| **UGT708A6** | 31.9 | 7.3 | NT | NT | NT |
| **UGT71A89** | NT | 34.7 | NT | NT | NT |
| **UGT71C1** | 98.5 | 98.9 | 100 | 63 | 26.6 |
| **UGT71D1** | 55 | 95.6 | 43.4 | NT | NT |
| **UGT71E1** | 32.6 | 22 | NT | NT | NT |

| | | | | | |
|---|---|---|---|---|---|
| **UGT71E5** | 70.3 | 10.7 | 47.4 | NT | NT |
| **UGT71F5** | 76.8 | 13.7 | 3.9 | NT | NT |
| **UGT72B10** | 97.3 | 8.3 | 3.3 | NT | NT |
| **UGT72B19** | 98.3 | 98.4 | 82.3 | 21.3 | 5 |
| **UGT72B68** | 95 | 97.9 | 98.3 | 46.2 | 0.0 |
| **UGT72B82** | 61 | 70.2 | 57.3 | NT | NT |
| **UGT72E2** | NT | 21.1 | NT | NT | NT |
| **UGT72E3** | NT | 14.5 | NT | NT | NT |
| **UGT72G4** | 40.0 | NT | NT | NT | NT |
| **UGT72J1** | 30.5 | NT | NT | NT | NT |
| **UGT74N7** | NT | 13.6 | NT | NT | NT |
| **UGT75L13** | NT | 11.2 | NT | NT | NT |
| **UGT75L5** | NT | 7.3 | NT | NT | NT |
| **UGT75L6** | NT | 0.0 | NT | NT | NT |
| **UGT78D2** | NT | 7.2 | NT | NT | NT |
| **UGT79B206** | NT | 8.2 | NT | NT | NT |
| **UGT85K6** | NT | 11.2 | NT | NT | NT |
| **UGT88A10** | 68.2 | 63.2 | 37.7 | NT | NT |
| **UGT88AE1** | 0.0 | 93.6 | 0.3 | NT | NT |
| **UGT88C1** | NT | 42.5 | NT | NT | NT |
| **UGT88C10** | 33.5 | 97.7 | 4.8 | NT | NT |
| **UGT88D7** | 0.0 | NT | NT | NT | NT |
| **UGT88E3** | 78.5 | 0.6 | 1.5 | NT | NT |
| **UGT88J1** | 49.1 | 3.9 | NT | NT | NT |

**Supplementary Table 6. UGT AEANT activity screening.** The % conversion yields from an overnight reaction between respective UGTs and substrates. NT, not tested.

| Impact category | Abbreviation | Unit | S1 | S2 | S3 |
|---|---|---|---|---|---|
| Freshwater ecotoxicity | FET | kg 1,4-DCB | 6.997 | 16.056 | 11.750 |
| Fine particulate matter formation | FPMF | kg PM2.5 eq | 0.437 | 0.737 | 0.511 |

| | | | | | |
|---|---|---|---|---|---|
| Marine eutrophication | ME | kg N eq | 1.383 | 1.176 | 1.078 |
| Ozone formation, Terrestrial ecosystems | OFT | kg NOx eq | 0.798 | 1.003 | 0.771 |
| Land use | LU | m2a crop eq | 1153.121 | 933.368 | 873.906 |
| Water consumption | WC | m3 | 20.470 | 20.208 | 16.834 |
| Ionizing radiation | IR | kBq Co-60 eq | 3.182 | 9.432 | 7.658 |
| Freshwater eutrophication | FE | kg P eq | 0.058 | 0.108 | 0.069 |
| Human non-carcinogenic toxicity | HNCT | kg 1,4-DCB | 312.785 | 433.102 | 258.173 |
| Marine ecotoxicity | MET | kg 1,4-DCB | 8.051 | 20.210 | 14.053 |
| Ozone formation, Human health | OFH | kg NOx eq | 0.779 | 0.979 | 0.754 |
| Terrestrial acidification | TA | kg SO2 eq | 1.647 | 2.671 | 1.942 |
| Mineral resource scarcity | MRS | kg Cu eq | 0.908 | 1.991 | 1.222 |
| Terrestrial ecotoxicity | TET | kg 1,4-DCB | 648.931 | 1583.210 | 1127.887 |
| Human carcinogenic toxicity | HCT | kg 1,4-DCB | 9.772 | 26.483 | 12.084 |
| Fossil resource scarcity | FRS | kg oil eq | 39.500 | 74.144 | 44.165 |
| Global warming | GW | kg CO2 eq | 248.085 | 401.165 | 245.856 |
| Stratospheric ozone depletion | OD | kg CFC11 eq | 0.002 | 0.003 | 0.002 |

**Supplementary Table 7**. **Impact on midpoint categories.** Midpoint categories for ReCiPe 2106 (H) and comparative analysis of three scenarios evaluated for the production of 100 kg of sunflower seeds. S1, Scenario 1; S2, Scenario 2; S3, Scenario 3.

| **Impact category** | **Abbreviation** | **Unit** | **S1** | **S2** | **S3** |
|---|---|---|---|---|---|
| Land use | LU | species.yr | 1.02E-05 | 8.29E-06 | 7.76E-06 |
| Global warming, Human health | GWH | DALY | 2.30E-04 | 3.72E-04 | 2.28E-04 |
| Marine ecotoxicity | MET | species.yr | 8.46E-10 | 2.12E-09 | 1.48E-09 |
| Fine particulate matter formation | FPMF | DALY | 2.75E-04 | 4.63E-04 | 3.21E-04 |
| Water consumption, Terrestrial ecosystem | WCT | species.yr | 2.76E-07 | 2.56E-07 | 2.27E-07 |

| | | | | | |
|---|---|---|---|---|---|
| Freshwater ecotoxicity | FET | species.yr | 4.85E-09 | 1.11E-08 | 8.14E-09 |
| Terrestrial ecotoxicity | TET | species.yr | 7.41E-09 | 1.81E-08 | 1.29E-08 |
| Water consumption, Human health | WCH | DALY | 4.54E-05 | 4.16E-05 | 3.74E-05 |
| Mineral resource scarcity | MRS | USD2013 | 2.10E-01 | 4.57E-01 | 2.83E-01 |
| Ozone formation, Terrestrial ecosystems | OFT | species.yr | 1.03E-07 | 1.29E-07 | 9.94E-08 |
| Global warming, Freshwater ecosystems | GWF | species.yr | 1.90E-11 | 3.07E-11 | 1.88E-11 |
| Human carcinogenic toxicity | HCT | DALY | 3.24E-05 | 8.79E-05 | 4.01E-05 |
| Fossil resource scarcity | FRS | USD2013 | 1.43E+01 | 2.48E+01 | 1.43E+01 |
| Freshwater eutrophication | FE | species.yr | 3.89E-08 | 7.21E-08 | 4.61E-08 |
| Human non-carcinogenic toxicity | HNCT | DALY | 7.13E-05 | 9.87E-05 | 5.89E-05 |
| Ozone formation, Human health | OFH | DALY | 7.09E-07 | 8.91E-07 | 6.87E-07 |
| Water consumption, Aquatic ecosystems | WCA | species.yr | 1.24E-11 | 1.57E-11 | 1.02E-11 |
| Marine eutrophication | ME | species.yr | 2.35E-09 | 2.00E-09 | 1.83E-09 |
| Ionizing radiation | IR | DALY | 2.70E-08 | 8.00E-08 | 6.50E-08 |
| Global warming, Terrestrial ecosystems | GWT | species.yr | 6.95E-07 | 1.12E-06 | 6.88E-07 |
| Terrestrial acidification | TA | species.yr | 3.49E-07 | 5.67E-07 | 4.12E-07 |
| Stratospheric ozone depletion | SOD | DALY | 1.14E-06 | 1.33E-06 | 9.86E-07 |

**Supplementary Table 8**. **Impact on endpoint categories.** Endpoint categories for ReCiPe 2106 (H) and comparative analysis of three scenarios evaluated for the production of 100 kg of sunflower seeds. S1, Scenario 1; S2, Scenario 2; S3, Scenario 3.

### Supplementary Figures

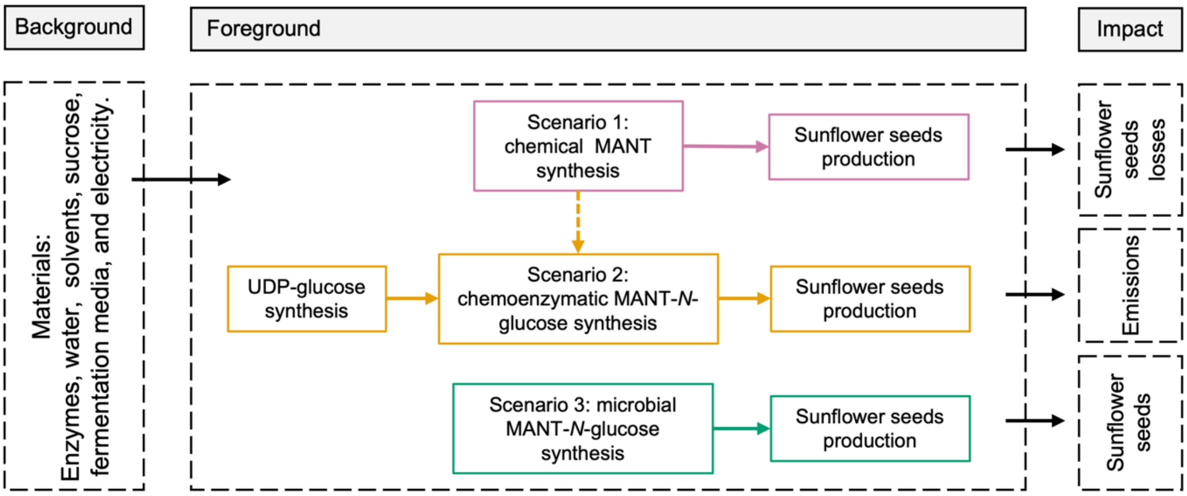


**Supplementary Figure 1. System boundaries and flow diagram of the preliminary LCA analysis.** The three scenarios compared are shown in pink (scenario 1), yellow (scenario 2), and green (scenario 3). The dashed arrow indicates that chemically synthesised MANT was used as the substrate for the chemoenzymatic synthesis of MANT-*N*-glucose.

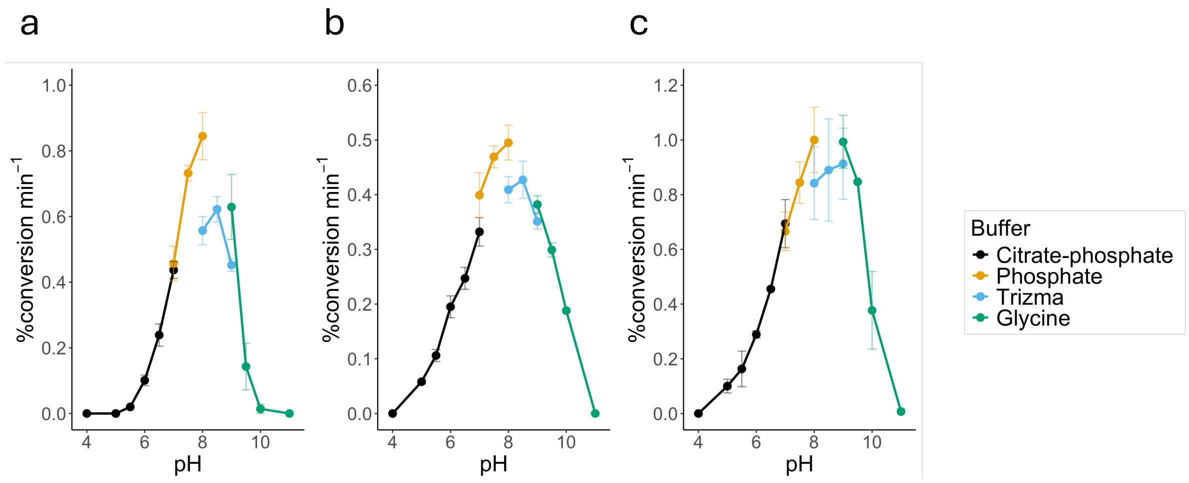


**Supplementary Figure 2. Optimal UGT reaction pH.** The optimal reaction pH of (a) UGT71C1, (b) UGT72B19, and (c) UGT72B68 using MANT as the acceptor substrate.

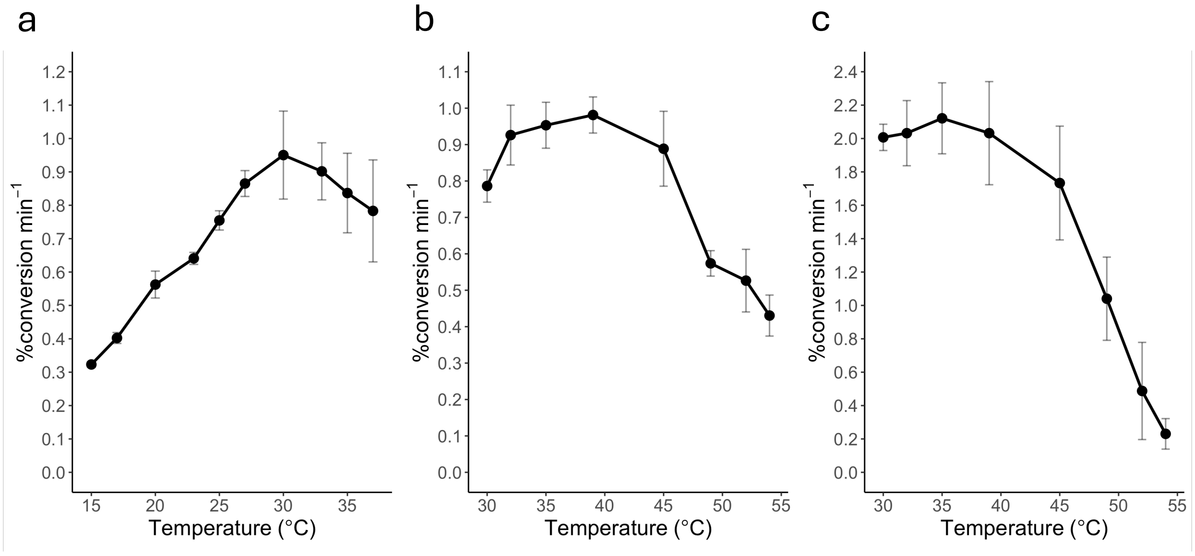


**Supplementary Figure 3. Optimal UGT reaction temperature.** The optimal reaction temperature of (a) UGT71C1, (b) UGT72B19, and (c) UGT72B68 using MANT as the acceptor substrate.

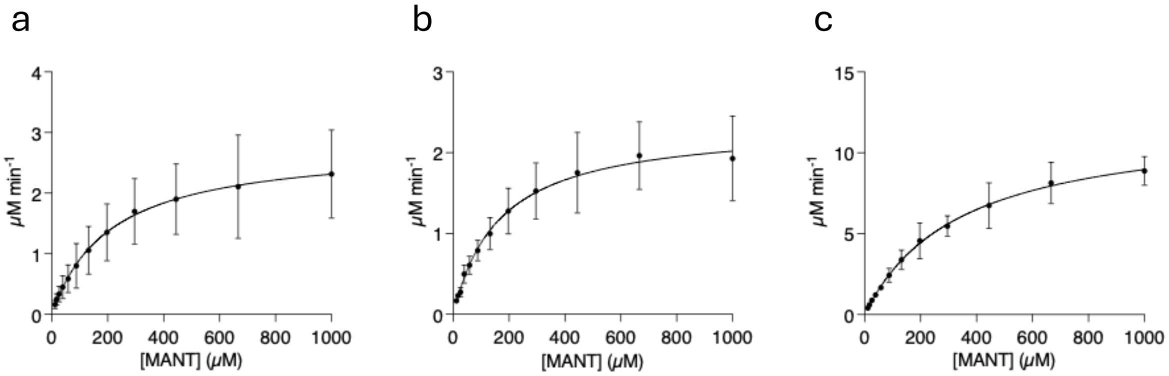


**Supplementary Figure 4. Kinetic characterisation, MANT.** The fitted Michaelis Menten plots of (a) UGT71C1, (b) UGT72B19, and (c) UGT72B68 using MANT as the acceptor substrate.

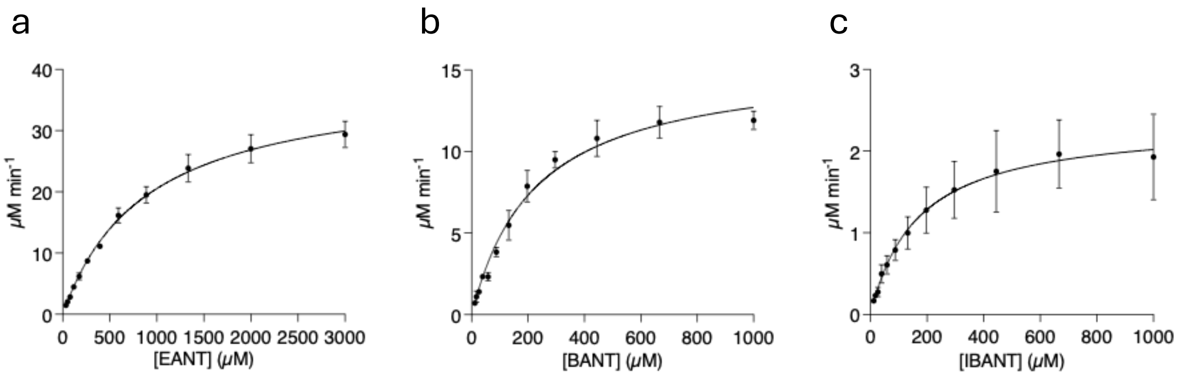


**Supplementary Figure 5. Kinetic characterisation, EANT, BANT, and IBANT.** The fitted Michaelis Menten plots of UGT72B68 using (a) EANT, (b) BANT, and (c) IBANT as the acceptor substrate.

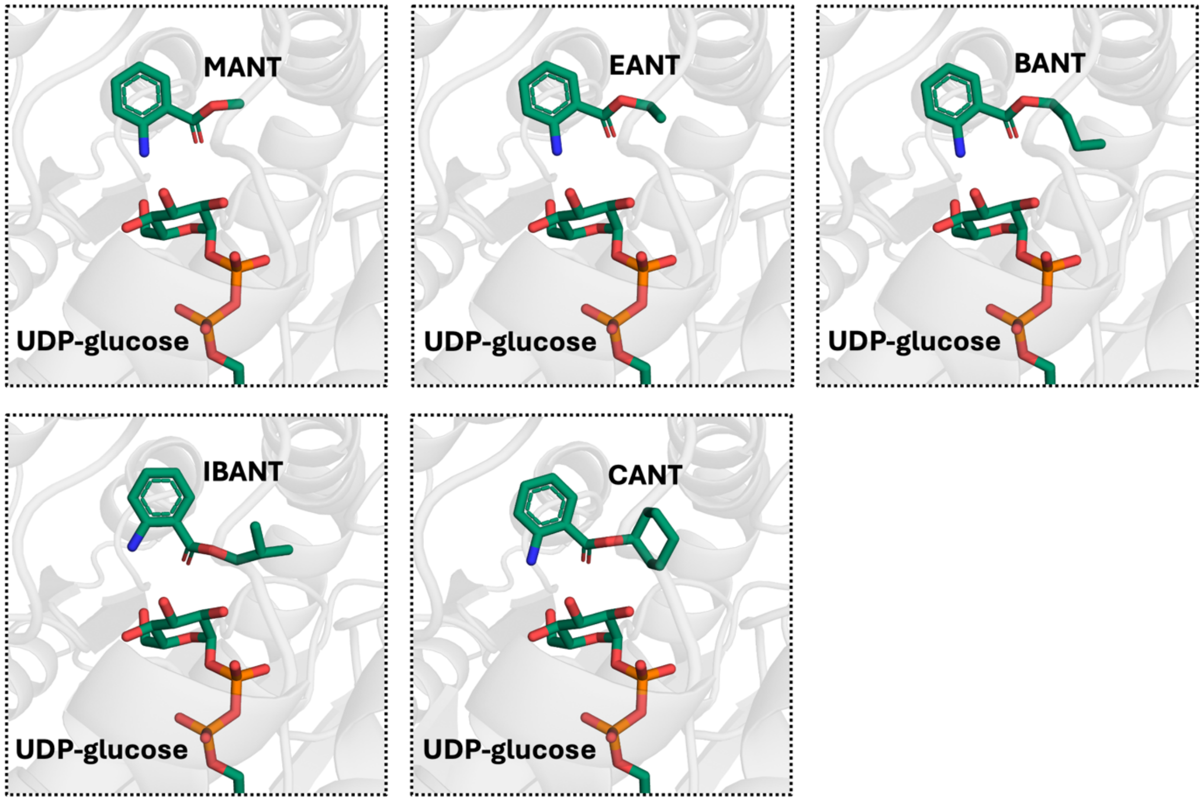


**Supplementary Figure 6. Molecular docking poses.** The docked poses of MANT, EANT, BANT, IBANT, and CANT in UGT72B68 with UDP-glucose superimposed into the model from PDB: 6SU6.[5]

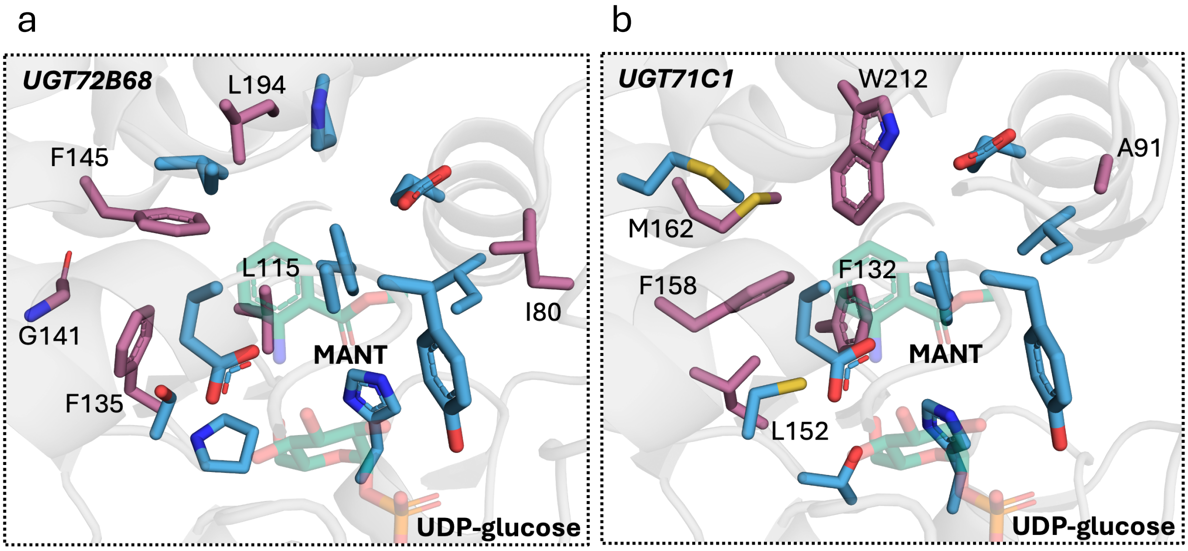


**Supplementary Figure 7. Active site residues, UGT72B68 and UGT71C1.** (a) MANT was docked into UGT72B68, and the residues within 5 Å of the docked MANT are shown as stick representations. Residues targeted for mutagenesis are shown in purple and labelled. (b) For UGT71C1, MANT binding coordinates were obtained by superimposing the docked MANT from UGT72B68 onto UGT71C1, and the corresponding residues within 5 Å of MANT are displayed as stick representations. The residues in UGT71C1 corresponding to the mutagenesis-targeted residues in UGT72B68 are shown in purple and labelled.

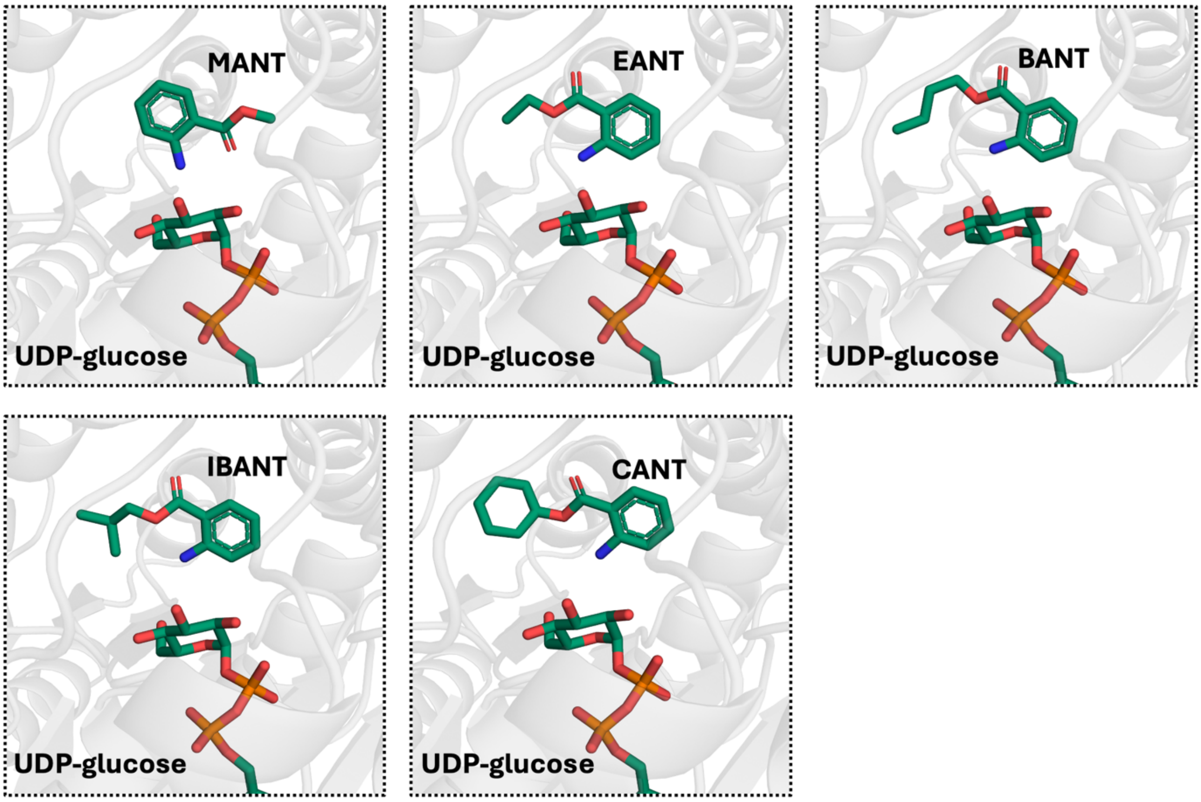


**Supplementary Figure 8. Chai-1 predicted binding poses.** The binding poses of MANT, EANT, BANT, IBANT, CANT, and UDP-glucose in UGT72B68 using the molecular structure prediction tool, Chai-1.[6]

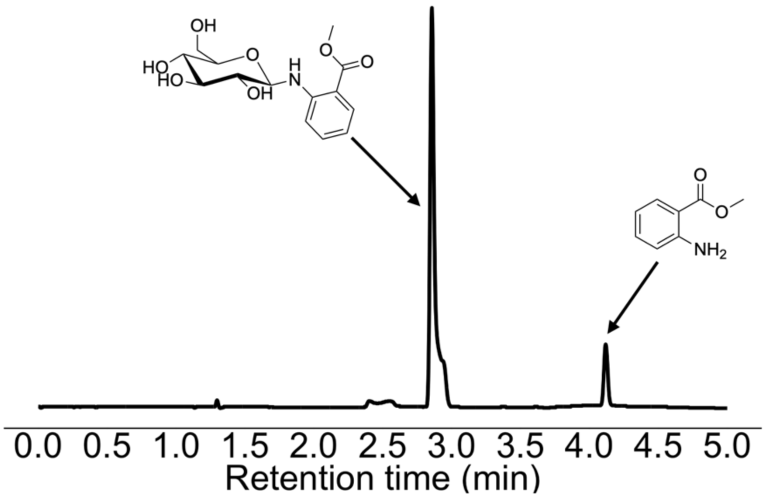


**Supplementary Figure 9. Biocatalytic MANT-*N*-glucose chromatogram.** HPLC chromatogram of the reaction mixture for the multigram-scale production after overnight incubation.

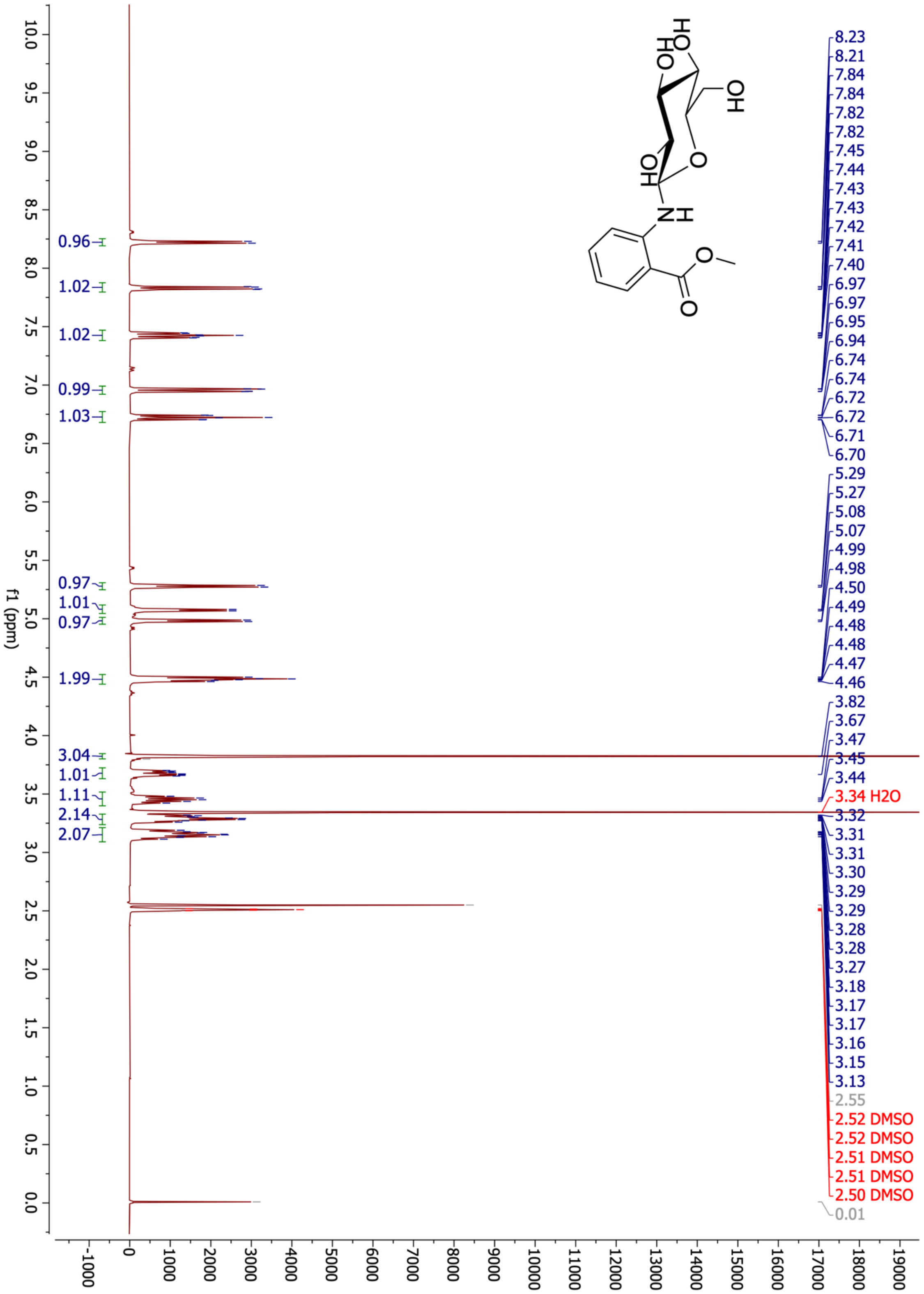


**Supplementary Figure 10. MANT-*N*-glucose H-NMR.** $^{1}$H NMR spectrum (400 MHz, DMSO-d6) of the isolated MANT-*N*-glucose: δ 8.22 (d, *J* = 6.2 Hz, 1H), 7.83 (dd, *J* = 8.1, 1.7 Hz, 1H), 7.43 (ddd, *J* = 8.7, 7.1, 1.7 Hz, 1H), 6.96 (dd, *J* = 8.7, 1.1 Hz, 1H), 6.72 (td, *J* = 7.6, 1.0 Hz, 1H), 5.28 (d, *J* = 5.9 Hz, 1H), 5.07 (d, *J* = 4.8 Hz, 1H), 4.98 (d, *J* = 5.2 Hz, 1H), 4.52 – 4.44 (m, 1H), 3.82 (s, 2H), 3.68 (ddd, *J* = 11.7, 5.3, 2.1 Hz, 1H), 3.45 (dt, *J* = 11.8, 5.9 Hz, 1H), 3.34 – 3.24 (m, 1H), 3.21 – 3.09 (m, 1H).

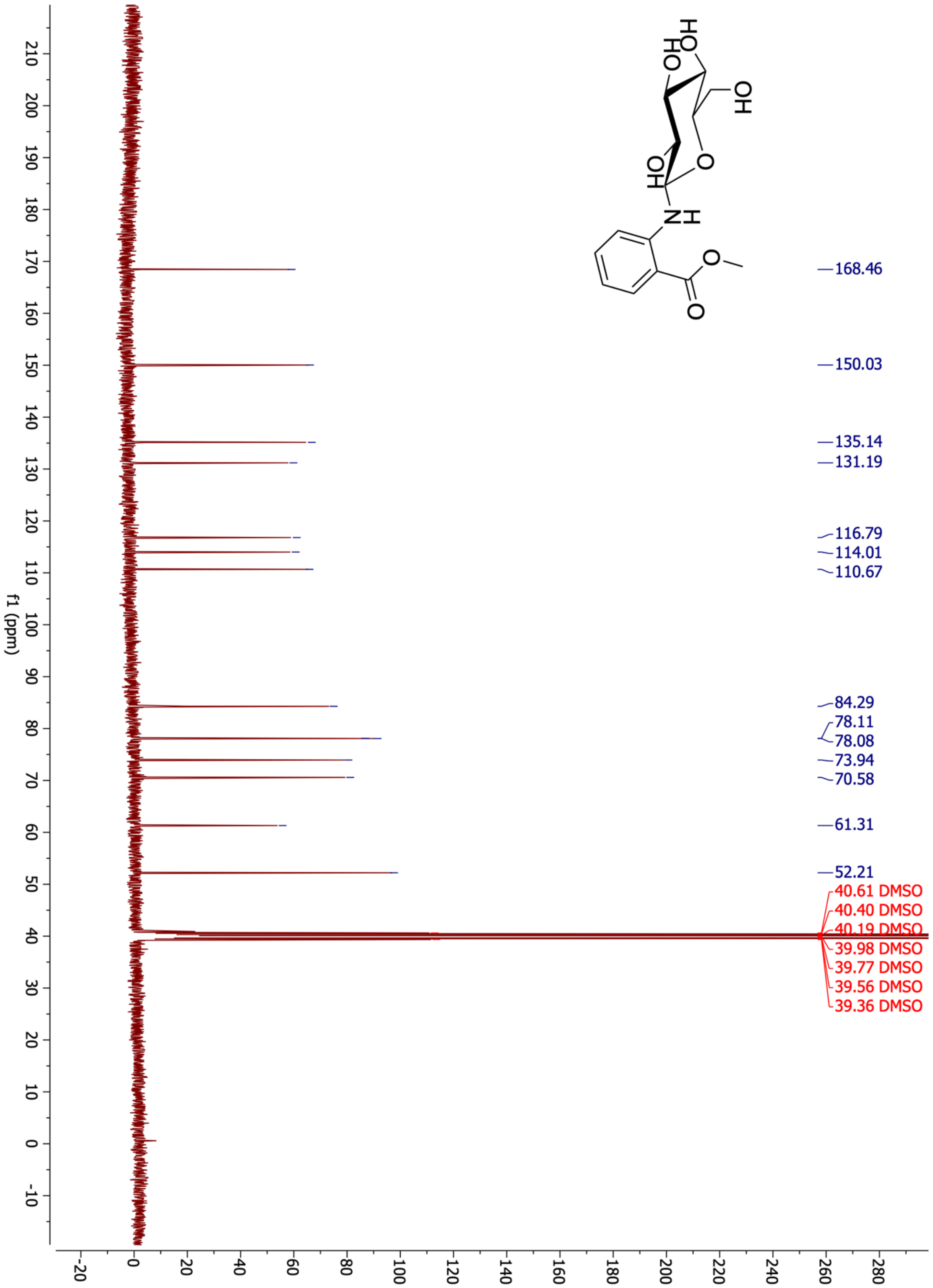


**Supplementary Figure 11. MANT-*N*-glucose C-NMR.** $^{13}$C NMR (101 MHz, DMSO-d6) of the isolated MANT-*N*-glucose: δ 168.46, 150.03, 135.14, 131.19, 116.79, 114.01, 110.67, 84.29, 78.11, 78.08, 73.94, 70.58, 61.31, 52.21

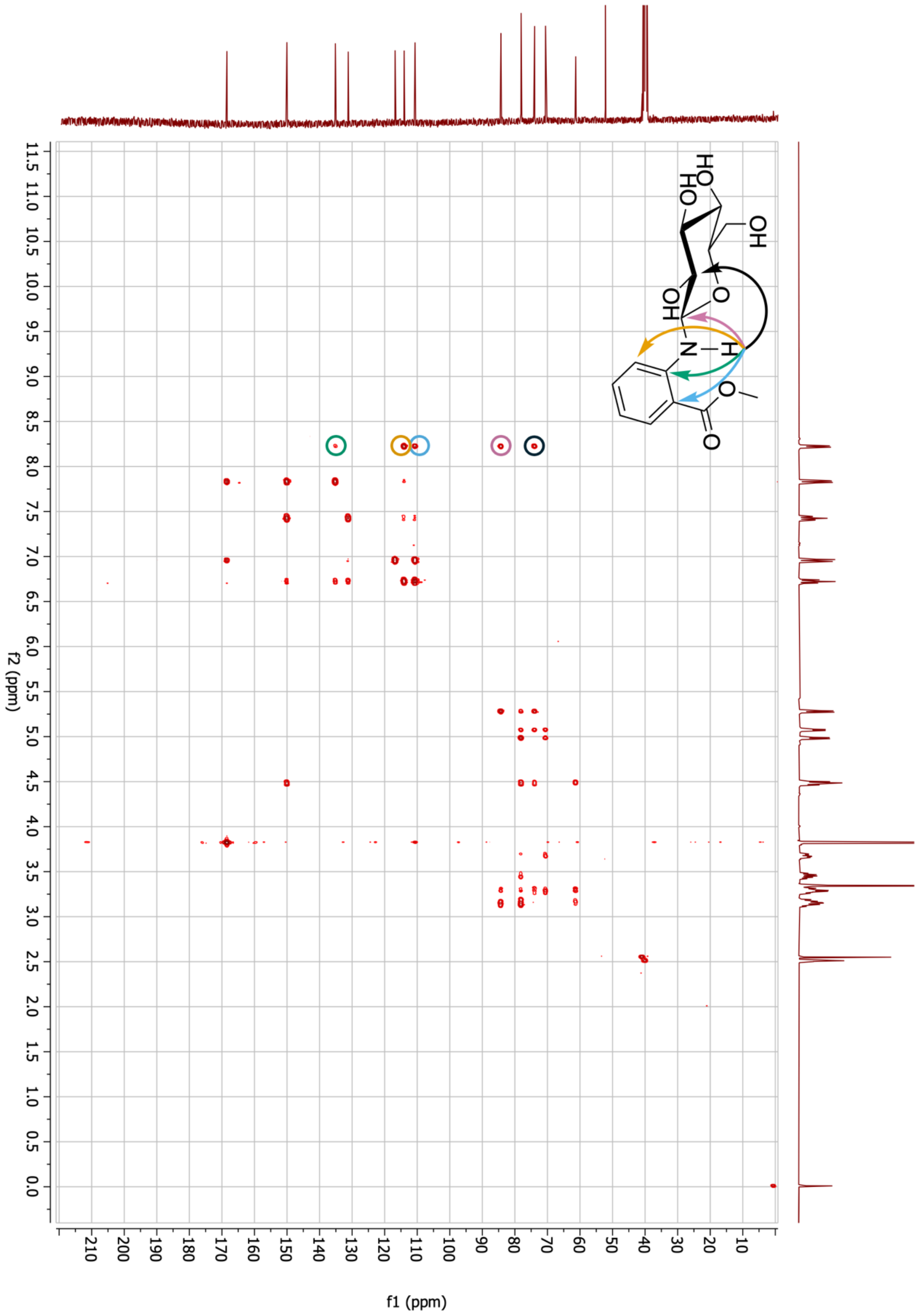


**Supplementary Figure 12. MANT-*N*-glucose HMBC.** HMBC spectrum of isolated MANT-N-glucose, highlighting signals corresponding to amine proton-carbon couplings. These signals are encircled in specific colours and used in the accompanying molecular model to indicate the exact proton-carbon couplings.

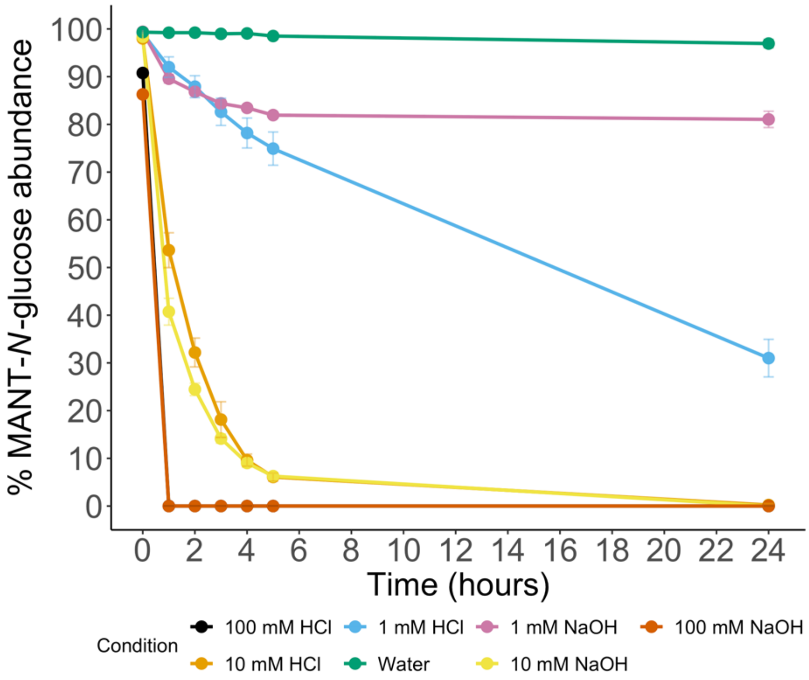


**Supplementary Figure 13. MANT-*N*-glucose chemical degradation.** The degradation of MANT-*N*-glucose in acidic, basic, and neutral conditions.

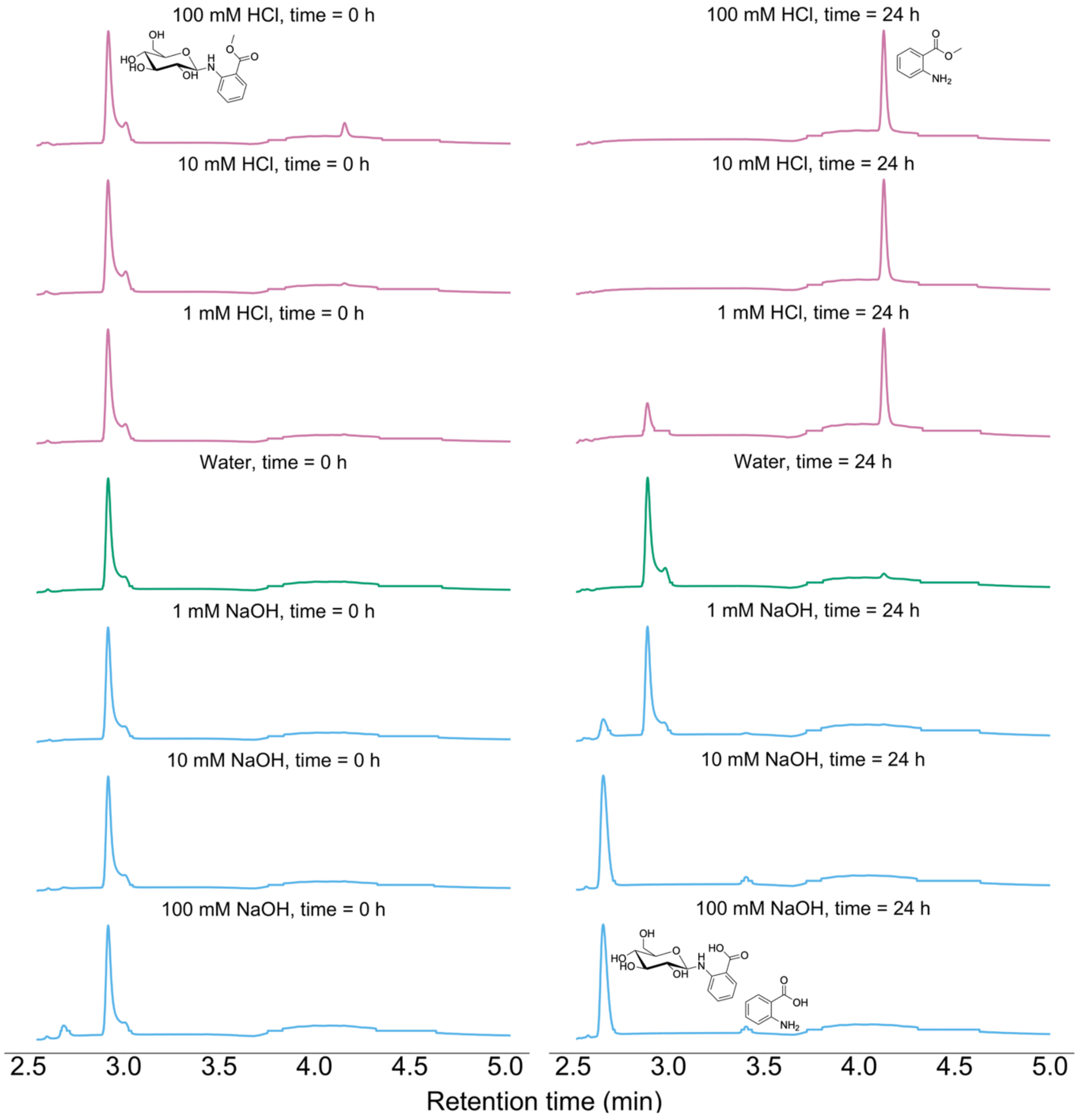


**Supplementary Figure 14. MANT-*N*-glucose chemical degradation products.** HPLC chromatograms of the degradation products of MANT-*N*-glucose at basic, acidic, and neutral conditions at time = 0 h and time = 24 h, respectively.

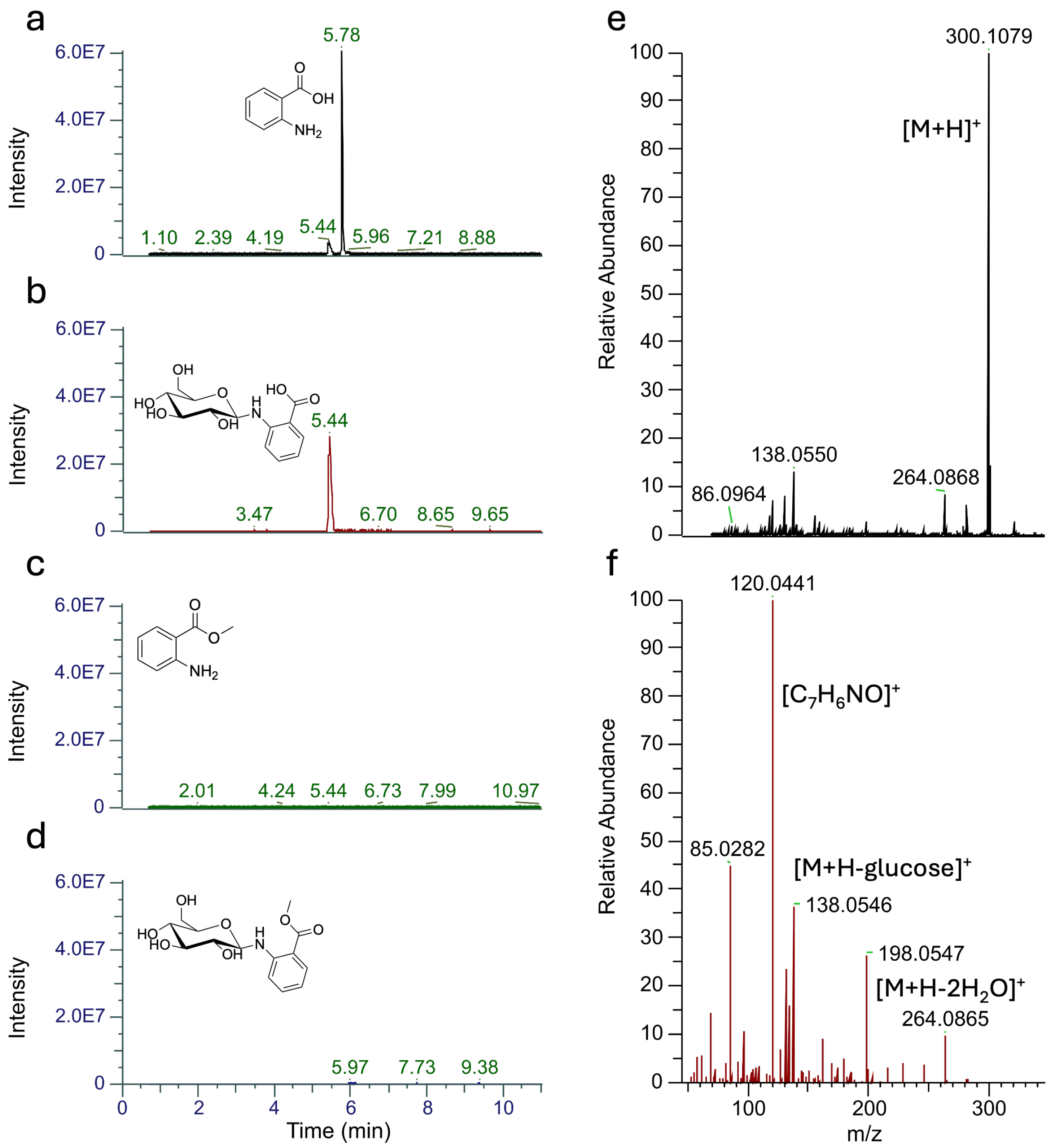


**Supplementary Figure 15. LC-MS validation of degradation product.** LC-MS chromatograms of the MANT-*N*-glucose in 100 mM NaOH reaction mixture after 24 hours. Ion traces at (a) m/z = 138.0536-138.0564 (ANT), (b) m/z = 300.1048-300.1108 (ANT-*N*-glucose), (c) m/z = 152.0689-152.0721 (MANT), and (d) m/z= 314.1202-314.1266 (MANT-*N*-glucose). (e) MS1 and (f) MS2 of the observed ion trace in (b) corresponding to ANT-*N*-glucose.

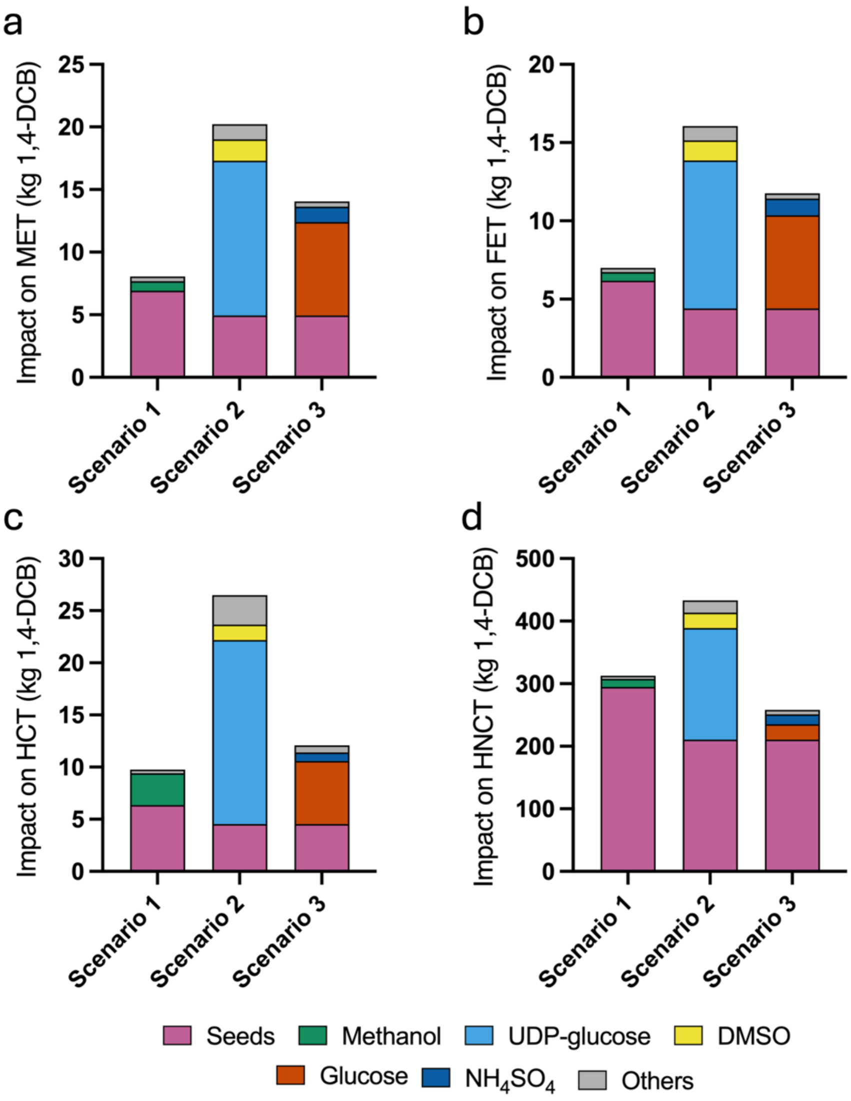


**Supplementary Figure 16. Contributors of the main impact categories.** Flow analysis of process factors in the main impact categories derived from the normalised analysis of midpoint categories: (a) MET, (b) FET, (c) HCT, and (d) HNCT. MET, marine ecotoxicity; FET, freshwater ecotoxicity; HCT, human carcinogenic toxicity; HNCT, human non-carcinogenic toxicity; 1,4-DCB, 1,4-dichlorobenzene.
